# Predicting the strength of urban-rural clines in a Mendelian polymorphism along a latitudinal gradient

**DOI:** 10.1101/635763

**Authors:** James S. Santangelo, Ken A. Thompson, Beata Cohan, Jibran Syed, Rob W. Ness, Marc T. J. Johnson

## Abstract

Cities are emerging as models for addressing the fundamental question of whether populations evolve in parallel to similar environments. Here, we examine the environmental factors that drive parallel evolutionary urban-rural clines in a Mendelian trait — the cyanogenic antiherbivore defense of white clover (*Trifolium repens*). We sampled over 700 urban and rural clover populations across 16 cities along a latitudinal transect in eastern North America. In each population, we quantified the frequency of genotypes that produce hydrogen cyanide (HCN), and in a subset of the cities we estimated the frequency of the alleles at the two genes (*CYP79D15* and *Li*) that epistatically interact to produce HCN. We then tested the hypothesis that winter environmental conditions cause the evolution of clines in HCN by comparing the strength of clines among cities located along a gradient of winter temperatures and frost exposure. Overall, half of the cities exhibited urban-rural clines in the frequency of HCN, whereby urban populations evolved lower HCN frequencies. The weakest clines in HCN occurred in cities with the lowest temperatures but greatest snowfall, supporting the hypothesis that snow buffers plants against winter frost and constrains the formation of clines. By contrast, the strongest clines occurred in the warmest cities where snow and frost are rare, suggesting that alternative selective agents are maintaining clines in warmer cities. Additionally, some clines were driven by evolution at only *CYP79D15*, consistent with stronger and more consistent selection on this locus than on *Li*. Together, our results demonstrate that both the agents and targets of selection vary across cities and highlight urban environments as large-scale models for disentangling the causes of parallel evolution in nature.

**Impact Summary:** Understanding whether independent populations evolve in the same way (i.e., in parallel) when subject to similar environments remains an important problem in evolutionary biology. Urban environments are a model for addressing the extent of parallel evolution in nature due to their convergent environments (e.g. heat islands, pollution, fragmentation), such that two distant cities are often more similar to one another than either is to nearby nonurban habitats. In this paper, we used white clover (*Trifolium repens*) as a model to study the drivers of parallel evolution in response to urbanization. We collected >11,000 plants from urban and rural habitats across 16 cities in eastern North America to examine how cities influence the evolution of a Mendelian polymorphism for an antiherbivore defense trait – hydrogen cyanide (HCN). This trait had previously been shown to exhibit adaptive evolution to winter temperature gradients at continental scales. Here we tested the hypothesis that winter environmental conditions cause changes in the frequency of HCN between urban and rural habitats. We found that half of all cities had lower frequency of HCN producing genotypes relative to rural habitats, demonstrating that cities drive parallel losses of HCN in eastern North America. We then used environmental data to understand why cities vary in the extent to which they drive reduction in HCN frequencies. The warmest cities showed the greatest reductions in HCN frequencies in urban habitats, while colder, snowier cities showed little change in HCN between urban and rural habitats. This suggests that snow weakens the strength of natural selection against HCN in cities. However, it additionally suggests alternative ecological or evolutionary mechanisms drive the strong differences in HCN between urban and rural habitats in the warmest cities. Overall, our work highlights urban environments as powerful, large-scale models for disentangling the causes of parallel and non-parallel evolution in nature.

## Introduction

The extent to which populations adapt in parallel to similar environmental conditions remains a fundamental problem in evolutionary biology (Losos 2017; Bolnick et al. 2018). High levels of genetic and phenotypic parallelism suggest that adaptive evolution is constrained, increasing our confidence in predicting species’ responses to similar conditions (Losos 2011). Despite predictions that similar environments should select for similar alleles and phenotypes, the degree of parallelism observed both within and among species is often imperfect (Bolnick et al. 2018). Genetic constraints, genetic drift, and gene flow, among other processes, can all alter the amount of parallelism among populations (Bolnick et al. 2018; Langerhans 2018). Replication is key to disentangling the many causes and consequences of parallel evolution in nature, from macroevolutionary (Schluter 2000) to microevolutionary scales (Lenski 2017), including leveraging naturally repeated cases of adaption across habitat types (Steiner et al. 2009; Stuart et al. 2017; Langerhans 2018).

Urban environments are emerging as models for investigating the causes of (non)parallel (*sensu* Bolnick et al. 2018) evolution among natural populations (Rivkin et al. 2019). Cities tend to share many biotic and abiotic environmental variables such as increased temperatures, elevated pollution, greater habitat fragmentation, and altered structure and composition of ecological communities (McKinney 2006), which can drive parallel adaptive evolution (Reid et al. 2016; Winchell et al. 2016; Yakub and Tiffin 2016; Kern and Langerhans 2018). Additionally, the commonly observed decreased size and increased isolation of urban populations can drive parallel losses of genetic diversity within urban populations due to stronger genetic drift and restricted gene flow (Munshi-South et al. 2016; Mueller et al. 2018). Despite the many examples of parallel responses to urbanization, imperfect parallelism—wherein phenotypes vary in the direction of evolutionary change between replicate urban and non-urban populations—is also common (Thompson et al. 2016; Diamond et al. 2018), although the causes of non-parallel responses to urbanization are poorly understood (Rivkin et al. 2019).

Recent work has used the globally-distributed plant, white clover (*Trifolium repens*) as a model for examining parallel evolutionary responses to urbanization. Thompson *et al.* (2016) documented repeated reductions in the frequency of HCN—a chemical plant defense against herbivores — within urban populations across three of the four cities examined in northeastern North America. Observational and experimental data show that HCN is predicted to be costly in the presence of cold winter temperatures because the metabolic components of HCN reduce tolerance to freezing (Daday 1954a, 1965; Kooyers et al. 2018). Consistent with this prediction, correlational data suggest that reduced urban snow cover has led to the observed colder winter ground temperatures in some cities relative to rural areas, which drives selection to reduce HCN frequencies in cities (Thompson et al. 2016). The absence of a cline in one of the four previously studied cities was explained by high urban snow depth in both urban and rural locations, which was hypothesized to insulate plants against the damaging effects of frost (Thompson et al. 2016). If urban-rural variation in snow depth is the only cause of urban-rural clines in cyanogenesis, this leads to an explicitly testable prediction: cities lacking snow should lack urban-rural clines in HCN. However, the two previous studies that have documented urban-rural cyanogenesis clines only sampled northern cities where minimum winter temperatures are below freezing (Thompson et al. 2016; Johnson et al. 2018), preventing a reliable test of the hypothesis that colder winter conditions are the primary agent driving the evolution of urban clines in HCN. Sampling cities that vary in winter temperature and frost exposure is required to understand the environmental conditions under which we expect to find (non)parallel responses of HCN to urbanization.

Because neutral processes can sometimes drive parallel phenotypic responses in nature (Losos 2011; Bolnick et al. 2018), it is important to reject neutral explanations before inferring that parallel selection has led to repeated adaptive differentiation. Population genetic simulations suggests that although drift could theoretically cause consistent clines if cities experience more drift (Santangelo et al. 2018a), empirical evidence from neutral microsatellite markers does not support this demography, implying selection is the primary driver of clines. (Johnson et al. 2018; Santangelo et al. 2018a), although three additional lines of evidence would help to distinguish between the roles of selection and drift in generating phenotypic clines in HCN. First, although the two-locus epistatic genetic architecture of HCN can lead to the evolution of clines due to drift (Santangelo et al. 2018a), neutral processes are expected to cause allele frequencies to vary randomly at the two underlying loci (Santangelo et al. 2018a). Thus, repeated clines in the same direction at individual loci underlying HCN can only be explained by selection driving differentiation of urban populations (Santangelo et al. 2018a). Second, an absence of clines in warm cities without snow would be consistent with altered selection in urban environments specifically caused by urban-rural gradients in snow depth and minimum winter ground temperatures. Additionally, clines in cyanogenesis to environmental gradients across North American have arisen via sorting of pre-existing and recurrent gene deletions at underlying loci (*CYP79D15* and *Li*) rather than novel mutations, such that multiple deletion haplotypes segregate at both loci in natural populations (Olsen et al. 2013; Kooyers and Olsen 2014; Olsen and Small 2018). The presence of multiple deletion haplotypes in both rural and urban populations would strengthen our inference that cyanogenesis is the target of selection rather than specific genes linked to the HCN loci; selection on linked genes are expected to lead to a single haplotype being overrepresented in urban or rural populations.

Here, we combine sampling of over 11,000 white clover plants from 16 cities with publicly available climate data to assess the environmental drivers underlying the evolution of latitudinal and urban-rural clines in cyanogenesis across multiple cities in eastern North America. We begin by assessing the environmental predictors of HCN frequencies along the latitudinal gradient by asking: (1) What regional environmental factors predict mean HCN frequencies within populations? Consistent with previous work (Daday 1954a, 1965, Kooyers and Olsen 2012, 2013), we expected to observe lower HCN frequencies at more northern latitudes due to lower winter temperatures. We then examine urban-rural clines in HCN across 16 cities to address the following questions: (2) How common are urban-rural clines among large (> 2 million people) cities in eastern North America? (3) What regional environmental factors predict the strength of clines in cyanogenesis? We predicted that we would observe the weakest clines in cities with high minimum winter temperature (i.e., warm cities) and also in those with high levels of snowfall due to weaker frost-mediated selection against HCN-producing genotypes. (4) Are clines present at both genes underlying cyanogenensis? Repeated clines at both loci underlying HCN would suggest that genetic drift does not cause urban-rural clines in HCN and that selection specifically acts on the production of HCN, as opposed to alternative functions of the individual loci. Finally, we ask: (5) Do urban populations show only a subset of the variation in deletion haplotypes as rural populations? The presence of multiple or fewer deletion haplotypes segregating in urban populations relative to rural populations would suggest that selection favors acyanogenic genotypes directly, rather than linked sites. Our results highlight urban environments as large-scale, replicated systems for addressing the ecological and genetic underpinnings of (non)parallel evolutionary responses in nature.

## Materials and methods

### Study system

*Trifolium repens* (Fabaceae) is a perennial legume that reproduces clonally through the production of stolons and sexually through self-incompatible, hermaphroditic flowers arranged in dense inflorescences (Burdon 1983). Plants are typically found in grazed or mowed pastures, lawns and meadows where they can maintain large dense populations (Burdon 1983). Native to Eurasia, *T. repens* was introduced to temperate regions worldwide as a forage and nitrogen-fixing crop (Burdon 1983; Kjærgaard 2003). Because of its long history of human-mediated transport, white clover is found in cities all over the world, making it an ideal system for studying patterns of parallel evolution in response to urbanization.

Many white clover populations are polymorphic for the production of hydrogen cyanide (HCN), with cyanogenic (HCN present) and acyanogenic (HCN absent) cyanotypes co-occurring (Daday 1958). The molecular and genetic basis at the individual loci underlying the production of both metabolic components involved in HCN production was recently characterized (Olsen et al. 2007, 2008, 2013; Olsen and Small 2018). The *Ac/ac* polymorphism is caused by deletions overlapping the *CYP79D15* locus (hereafter *Ac*), which encodes the cytochrome P450 subunit involved in the synthesis of cyanogenic glycosides (linamarin and lotaustralin) stored in the cell vacuole (Olsen et al. 2008, 2013; Olsen and Small 2018). Plants require at least one functional allele with dominant expression (i.e., *Ac*–) to produce cyanogenic glycosides. Similarly, the *Li/li* polymorphism results from a deletion at the *Li* locus encoding the hydrolyzing enzyme linamarase, which is stored in the cell wall (Kakes 1985); at least one dominant allele (i.e., *Li*–) is required to produce linamarase. Thus, plants require a minimum of one dominant allele at each locus to produce HCN (i.e., cyanotype *Ac*– *Li*–), which is released when cell damage causes cyanogenic glycosides and linamarase to interact (Hughes 1991). If either locus is homozygous for the recessive allele, then a plant lacks HCN and is said to be “acyanogenic” (i.e., cyanotypes *Ac*– *lili, acac Li*–, *acac lili*).

### Sampling and HCN assays

In May and June 2016, we sampled 15 plants from each of 15 to 45 populations (mean = ∼38) along urban-rural gradients in each of 12 cities in eastern U.S.A (Fig. 1). Power analyses conducted by resampling the data from Thompson *et al.* (2016) showed that this sampling scheme provides sufficient power to detect even the weakest statistically significant clines in HCN (see supplemental text S2: “Power analyses for sampling design”, Fig. S1). We sampled only large cities (240,000 < Human population size (city area) < 8,200,000; 151 < city area (km^2^) < 2,300.) because these are likely to have the strongest environmental gradients associated with urbanization. We additionally chose cities along a north-south latitudinal transect such that more southern cities had less snow and warmer winter ground temperatures, which earlier research suggested would weaken selection against HCN in urban environments, leading to weaker or absent clines in southern cities (Thompson et al. 2016).

**Figure 1:**
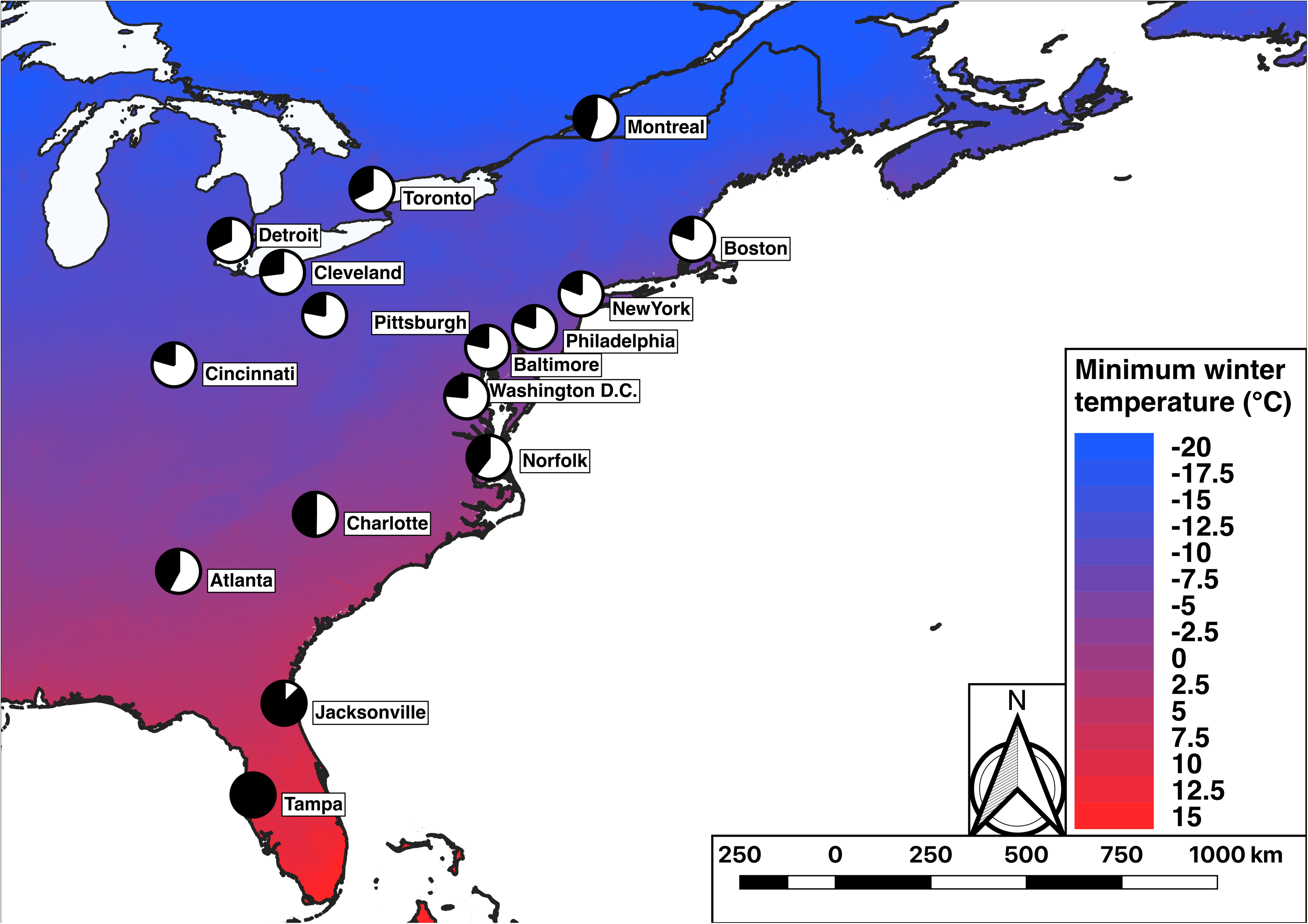
Map of 16 cities from which we sampled white clover populations along urban-rural transects. Pie charts represent the mean frequency of HCN (black = HCN+, white = HCN−) for each city when averaged across all populations along the transect. Map color depicts the gradient in the minimum winter temperature (MWT, °C) taken from BioClim.

Sampling took place in three trips: trip one (May 16^th^ to 23^rd^, 2016) involved collections from Tampa and Jacksonville, FL, Atlanta, GA, and Charlotte, NC; trip 2 (June 5^th^ to 11^th^, 2016) from Norfolk, VA, Washington, D.C., and Baltimore, MD, and Philadelphia, PA; and trip 3 (June 15^th^ to 20^th^) from Cleveland and Cincinnati, OH, Pittsburgh, PA, and Detroit, MI. In each city, we targeted populations spaced at least 1 km apart. In each population, we recorded the latitude and longitude coordinates and collected ∼6 cm-long white clover stolons with three to four intact leaves; stolons were at least 1.5 m apart to minimize sampling the same clonal genotype. Stolons were placed in sandwich bags and kept on ice in a cooler until being brought back to the lab where they were individually placed in 2 ml microcentrifuge tubes and stored at −80 °C until HCN phenotyping. In total, we collected and assayed 6,738 stolons from 459 populations across 12 cities. For all downstream analyses, we combined the data from the 12 cities described above with the 4 cities (Boston, MA, New York, NY, Toronto and Montréal, CA) originally sampled by Thompson *et al.* (2016). In total, we analyzed urban-rural clines in HCN using 11,908 plants from 721 populations across 16 cities.

We used Feigl-Anger assays to determine whether plants were cyanogenic or acyanogenic (Feigl and Anger 1966; Gleadow et al. 2011), which uses a simple color change reaction to determine the plant’s phenotype. Briefly, we added a single mature leaf to wells in 96-well plates with 80 μL of sterile deionized water. Leaf samples were added to alternating wells so that a single plate could accommodate 48 plant samples. The plates were frozen at −80°C to facilitate cell-lysis and release of HCN, and upon removal we macerated the plant tissue with pipette tips. We then covered the plate with Feigl-Anger test paper and incubated the covered plate at 37°C for 3 hr. Cyanogenic individuals (i.e., *Ac- Li-*) produce a blue spot on the filter paper above the well, whereas acyanogenic plants (i.e., *Ac- lili, acac Li-*, or *acac lili*) produce no color change.

To assess whether clines were driven by changes in the frequency of HCN or clines at either component gene (i.e. *Ac* or *Li*), we determined the frequency of *Ac* and *Li* for a subset of the cities (Atlanta, Baltimore, Charlotte, Cleveland, Jacksonville, New York, Norfolk, and Washington), which was combined with previously-collected allele frequency information for *Ac* and *Li* for the city of Toronto (Thompson et al. 2016). For plants that tested negative for HCN, we added either (1) 30 μL of 10 mM exogenous cyanogenic glycosides (linamarin, Sigma-Aldrich 68264) plus 50 uL of ddH_2_0 or (2) 80 μL of 0.2 EU / mL linamarase (LGC Standards CDX-00012238-100). A positive reaction in (1) indicates a plant producing linamarase (i.e. *acac Li*–); a positive reaction in (2) indicates a plant producing glycosides (i.e. *Ac*– *lili*); a negative reaction in both indicates plants that do not produce glycosides nor linamarase (i.e. the double-homozygous recessive genotype, *acac lili*). These assays have been previously confirmed through PCR to reliably determine the cyanotype of individual *T. repens* genotypes (Olsen et al. 2007, 2008; Thompson and Johnson 2016). Due to the complete dominance of functional alleles at both loci, we are unable to calculate the frequency of *Ac* or *Li* solely from the phenotyping assays described above (e.g. *AcAc* and *Acac* are indistinguishable). We therefore used the marker frequency (e.g. *Ac*–) to calculate the frequency of *Ac* and *Li* assuming Hardy-Weinberg equilibrium (*p*^2^ + 2*pq* + *q*^2^ = 1), where *q*^2^ represented the observed frequency of homozygous recessive genotypes at *Ac* (i.e. *acac*) or *Li* (i.e. *lili*). While urban-rural HCN clines may not always meet the assumptions of HWE (Johnson et al. 2018; Santangelo et al. 2018a), deviations from HWE are not predicted to greatly impact inferred allele frequencies when homozygous dominant and heterozygous individuals are phenotypically identical, as is the case for HCN (Lachance 2009; Kooyers and Olsen 2013). Analyzing changes in the frequency of *Ac* and *Li* for these nine cities allowed us to assess whether selection is targeting HCN specifically or individual loci underlying HCN production.

To examine whether urban and rural populations varied in the frequency of deletion haplotypes at *Ac* or *Li*, we used PCR with previously described forward-reverse primer pairs (Olsen et al. 2014) to identify the relative size of deletions at both loci for individual plants. We extracted total genomic DNA from each of 10 randomly-selected urban and rural plants (n = 20) for each of seven cities (Table 1) using a standard CTAB-chloroform extraction method (Agrawal et al. 2012). We chose these cities because they spanned the range of latitudes included in our study, thus reducing potential impact of geographical variation on haplotype frequencies. We included cities that varied in the presence (5 cities) and absence (2 cities) of clines in HCN. We only extracted DNA from plants that were homozygous recessive at both loci (i.e. *acac lili*) because these plants have at least one deletion haplotype at each locus. Each plant was amplified with 6 different primer pairs (3 for each locus), designed to assay the approximate size of the genomic deletion at each locus based on the presence/absence of PCR products (Kooyers and Olsen 2014). Note that larger deletions are masked by smaller deletions when resolving haplotypes on a gel, preventing us from estimating the true frequency of each deletion; we therefore rely only the presence/absence of deletions in our analyses (see “Statistical analyses” below). Using this approach, we were able to assign 88% and 78% of samples to previously described *Li* (n = 4) and *Ac* (n = 2) deletion haplotypes, respectively. The remaining individuals were either newly discovered haplotypes or individuals with intact *Li* and *Ac* genes, the latter indicating either false negative phenotyping or false positive haplotyping assays (or the presence of a silencer modifier locus). We focus our results on the haplotypes that aligned with those previously described by Kooyers and Olsen (2014) to understand whether one deletion haplotype versus multiple haplotypes were segregating in urban and rural populations in cities across our latitudinal transect.

**Table 1.**
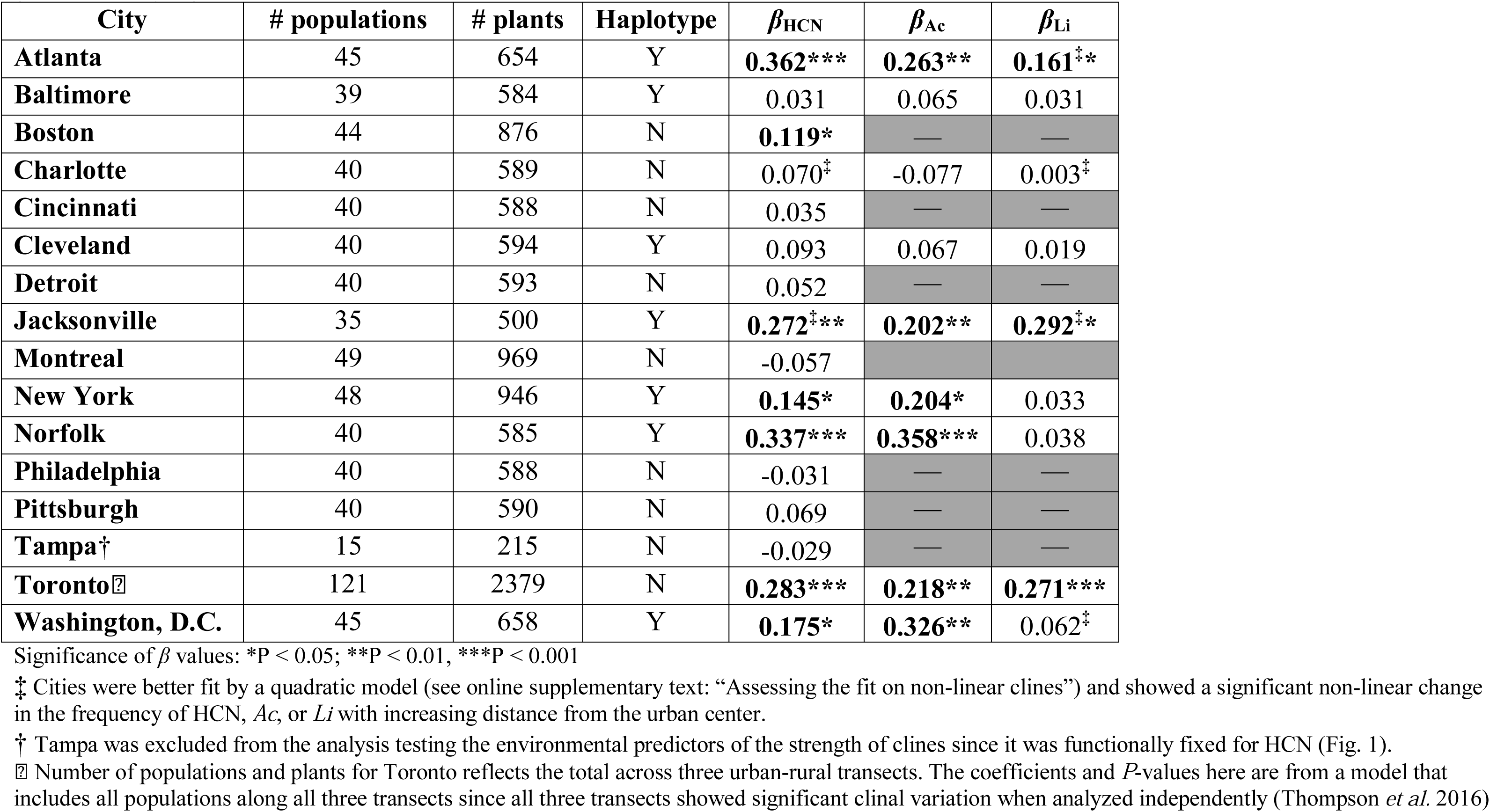
Beta coefficients (i.e. slope) and *P-*values from linear models testing the change in the frequency of HCN, *Ac*, or *Li* with increasing distance (standardized) from the urban center for each of 16 cities. Also shown are the total number of populations and plants sampled in each city, and whether deletion haplotypes were identified in urban and rural population of that city. Bolded terms represent linear clines that were significant at *P* < 0.05. Grey boxes represent cities where we did not quantify the frequency at the genes underlying HCN.

### Environmental data

To examine the regional abiotic factors that predict the strength of clines in HCN, we retrieved environmental data from publicly available databases. First, we retrieved the minimum winter temperature (MWT, Bio6) — an important predictor of HCN frequencies (Daday 1965; Kooyers and Olsen 2013) — and the maximum summer temperature (MST, Bio5) using the highest resolution data (30 arc seconds; 1 km^2^) available from the BioClim database (version 1.4, Hijmans et al. 2005). We additionally retrieved the average monthly precipitation (Precip) at 1 km^2^ resolution from the same database. Next, we obtained the annual aridity index (AI), monthly average potential evapotranspiration (mPET) and average annual potential evapotranspiration (aPET) at 1 km^2^ resolution from the Consortium for Spatial Information (CGIAR-CSI, Trabucco and Zomer 2009). These three abiotic factors are known predictors of the frequency of HCN and its component genes at the continental scale in North America, Europe and New Zealand (Kooyers and Olsen 2013). BioClim and CGIAR datasets are provided as gridded raster layers, from which we extracted the relevant data for all 721 populations using QGIS v. 3.2.3 (QGIS Development Team 2018).

Finally, we obtained daily snow depth, snowfall, maximum temperature, and minimum temperature for all cities for the years 1980 to 2015 from the National Oceanic and Atmospheric Administration’s National Centers for Environmental Information (https://www.ncdc.noaa.gov/cdo-web/datatools/selectlocation). Importantly, these are regional environmental data obtained from a single weather station for each city located at the nearest international airport; thus, these data represent city-level abiotic conditions and not the conditions extracted for each population.

Some filtering and processing of the environmental data was required prior to downstream analyses. First, we took the mean MWT (°C), MST (°C), AI, and aPET across all populations within a city to estimate the city-level minimum winter temperature, maximum summer temperature, aridity, and annual potential evapotranspiration, respectively. Next, we calculated an alternative measure of aridity, the soil moisture deficit (SMD), as the difference between Precip and mPET. This measure of aridity can more reliably estimate regional aridity in cases where both precipitation and PET are low (Thompson et al. 2014). We calculated SMD for all months from May to September to represent the plant growing season, and again took the mean of these measurements across all populations as our measure of city-level SMD. Finally, we used NOAA’s weather station data to estimate regional snow depth (cm), snowfall (cm), and the number of days below zero with no snow cover, a measure of frost exposure that has been previously associated with urban-rural clines in HCN (Thompson et al. 2016). To retrieve these estimates, we first filtered the weather data to remove missing data and only included observations from January and February as these are the coldest winter months in eastern North America. We additionally removed observations from years where data was unavailable for both January and February and eliminated months with fewer than 10 days of weather data. Following filtering, we retained 31,005 weather observations, with a mean of 1937 observations per city (Table S1). From these data, we calculated the mean snow depth, mean snowfall, and the number of days below 0 °C with no snow cover (i.e. snow depth of 0 cm) and took the mean of these measurements across all years as our estimates of regional winter conditions.

### Statistical analyses

For brevity, we only briefly describe the statistical procedures used throughout the paper; a detailed description of all analyses can be found in the supplementary materials (text S1: “Detailed statistical analyses”). We first tested whether, on average, cities varied in mean HCN frequencies and whether urbanization influenced HCN frequencies. To do this, we fit an ANOVA using type III SS with within-population HCN frequencies as the response variable and city, standardized distance to the urban center and the city × distance interaction as predictors. We used distance to the urban center as a measure of urbanization as this is highly correlated with % impervious surface (R^2^ = 0.64, Johnson et al. 2018) and sufficiently captures variation in HCN frequencies across urban-rural gradients (Thompson et al. 2016; Johnson et al. 2018). Because urban-rural transects varied in length, we scaled distance within cities between 0 (most urban) and 1 (most rural). In our model, a significant effect of City suggests that mean HCN frequencies vary across the 16 cities. A significant Distance term means that across all cities, HCN frequencies vary in parallel across the urban-rural transect (i.e. parallel clines in HCN frequencies), while a significant City × Distance interaction indicates the strength or direction of clines in HCN varies across cities. The significant effects of City and the City × Distance interaction in our model (see “Results”) justify an examination of the environmental predictors of mean HCN frequencies and variation in the strength of clines across cities, respectively.

To assess the environmental predictors of mean HCN frequencies across cities along our latitudinal gradient, we fit the following linear model: mean HCN frequency ∼ PC1_HCN_ + # days < 0°C with no snow + annual aridity index + soil moisture deficit, where PC1_HCN_ is a composite axis generated via PCA that explained 90.2% of the variation in MST, MWT, summer precipitation, annual PET, and snowfall, all of which were highly correlated and individually significantly predicted variation in HCN frequencies. Lower values of PC1_HCN_ represented cities with higher summer temperatures, higher minimum winter temperatures, higher summer precipitation, greater potential evapotranspiration, and lower snowfall. We assessed significance of model predictors using an AIC_c_-based multi-model selection and averaging process.

Upon confirming that cities varied significantly in the strength of urban-rural phenotypic clines using ANOVA, we used a similar approach to that described above to examine the environmental predictors of the strength of urban-rural phenotypic clines in HCN. For each city, we first fit a linear regression with the proportion of cyanogenic plants within each population as the response variable and standardized distance to the urban center as the sole predictor. Note that cities that showed significant changes in HCN frequency with distance (Table 1) were also significant following Bonferroni correction of logistic regressions using data from individual plants (i.e., 1 for HCN+, 0 for HCN−). We extracted the slope (i.e. *β* coefficient) from each city’s model as a measure of the strength of the clines and examined the environmental predictors of cline strength by running the following model: *β* ∼ PC1_slope_, where PC1_slope_ is a composite axis generated via PCA that explains 92.8% of the variation in snow depth, snowfall, MWT, and MST, all of which were highly correlated and on their own significantly predicted variation in the strength of clines (see text S1). Cities with low values along PC1_slope_ experience little snow, higher minimum winter temperature, and higher maximum summer temperature.

We explored whether clines were present at each of the two loci underlying HCN. This was done by fitting linear models in which the allele frequencies for the *Ac* and *Li* loci were treated individually as a response variable in separate analyses, which was regressed against standardized distance to the urban core as a predictor. Finally, to examine variation in deletion haplotypes across urban and rural habitats, we used the raw counts of deletion haplotypes at each locus to calculate Simpson’s diversity index for deletions in urban and rural habitats for each city, which accounts for both presence/absence and abundance of haplotypes in each population when estimating diversity. We fit Simpson’s diversity index as the response variable in a linear model with habitat type (i.e. urban vs. rural) as the sole predictor such that a significant effect of habitat suggested differences in deletion haplotype diversity among urban and rural habitats. All analyses were performed in R v. 3.6.0 (R Core Team 2019).

## Results

### Variation in HCN frequencies in cities along a latitudinal gradient

The mean frequency of HCN varied across the 16 cities from 19% (New York) to 99% (Tampa) (Effect of City: F_15,689_ = 18.48, *P* < 0.001, Table 1, Fig. 1, Fig. 2), with the highest frequencies at the most southern latitudes (Fig. S2). The number of days < 0°C with no snow cover and PC1_HCN_ together accounted for 94.1% of the variation in mean HCN frequencies among cities (Table S4). Specifically, HCN frequencies decreased by 1.5% for every additional day below 0 °C with no snow cover (*β* = −0.015, *Z* = 8.77, *P* < 0.001, Table S5, Fig. 3a), and by 6.4% for every unit increase along PC1_HCN_ (*β* = −0.064, *Z* = 7.0, *P* < 0.001, Table S5, Fig. 3b), suggesting HCN frequencies decrease in colder, wetter environments that get more snow. Annual aridity index was not significant predictor of mean HCN frequencies in our model (*P* = 0.36, Table S5) while soil moisture deficit was not included in any top models following model selection and averaging.

**Figure 2:**
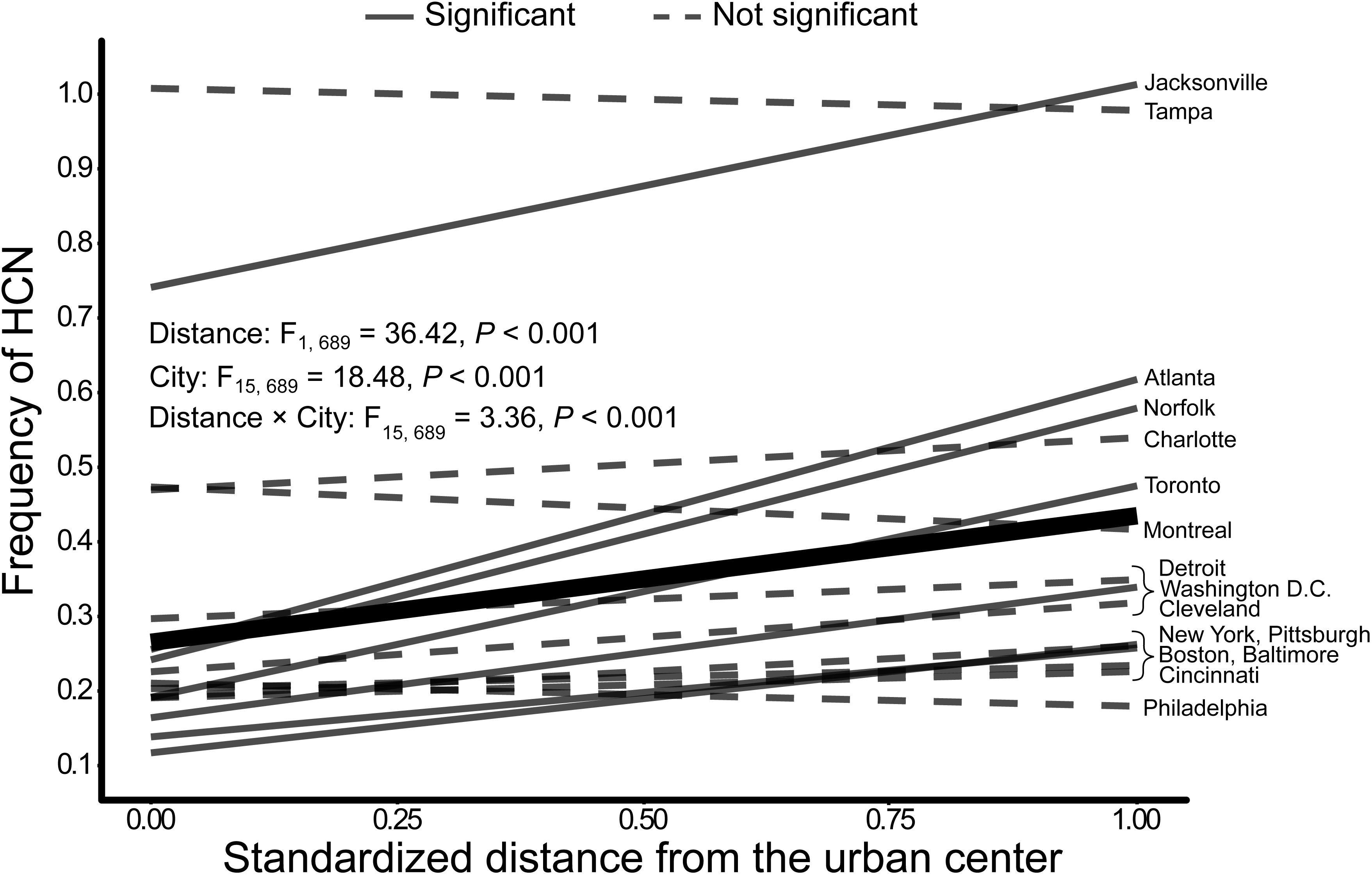
Urban-rural clines in the frequency of HCN within populations of *Trifolium repens* across 16 cities in eastern North America. The frequency of HCN within *T. repens* population is plotted against the standardized distance from the urban center. Solid lines represent linear regressions from cities where the phenotypic cline in HCN was significant at *P* < 0.05, whereas dashed lines are cities that lack significant clinal variation. The thick black line represents the main effect of standardized distance on HCN frequencies, averaged across all cities.

**Figure 3:**
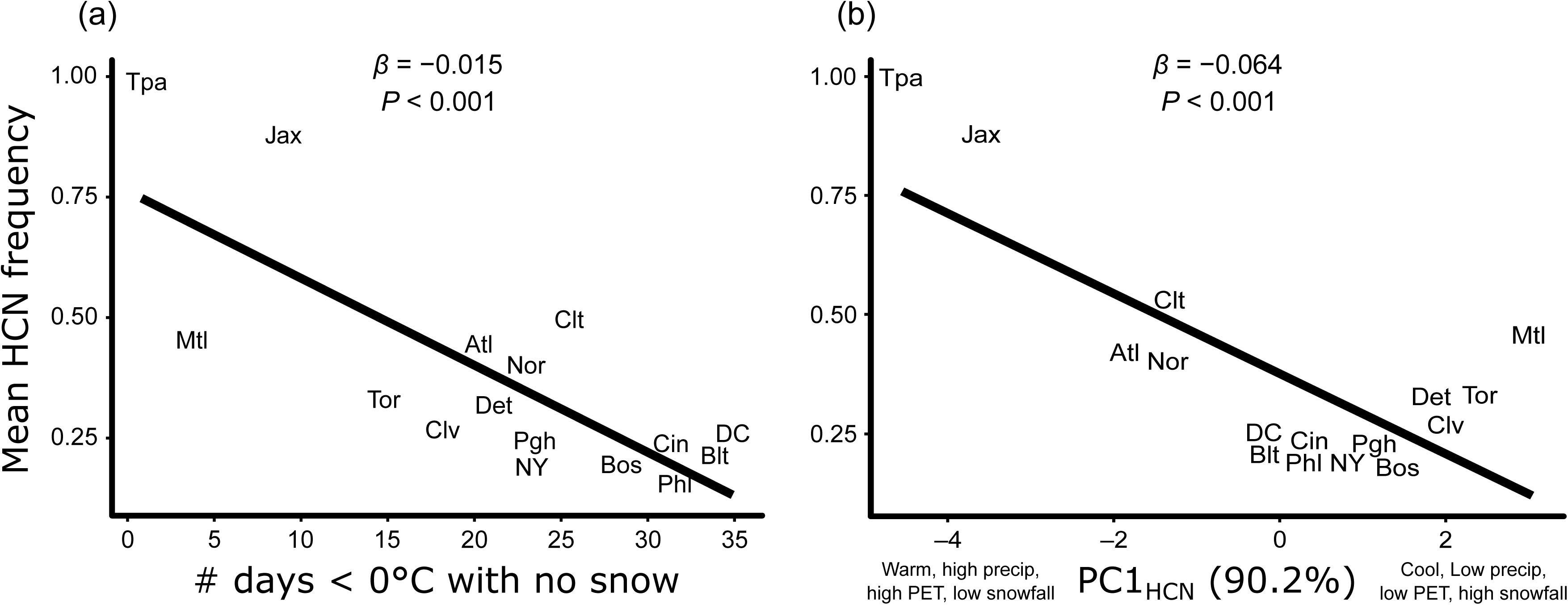
Mean HCN frequency was influenced by (a) the number of days below 0 °C with no snow cover — a measure of frost exposure — and (b) PC1_HCN_, a component axis accounting for 90.2 % of the variation in maximum summer temperature (°C, Bio5), minimum winter temperature (°C, Bio6), annual potential evapotranspiration (mm), monthly summer precipitation (mm), and snowfall (cm) (inset in (b)). City labels are slightly jittered to avoid overlap, if necessary. Cities with low values along PC1_HCN_ have high summer temperatures, high minimum winter temperatures, high summer precipitation and potential evapotranspiration, and low snowfall, whereas cities with high values along PC1_HCN_ have the opposite. (City abbreviations: Jacksonville (Jax); Tampa (Tpa); Atlanta (Atl); Norfolk (Nor); Charlotte (Clt); Toronto (Tor); Montréal (Mtl); Detroit (Det); Washington D.C. (DC); Cleveland (Clv); New York (NY); Pittsburgh (Pgh); Boston (Bos); Baltimore (Blt); Cincinnati (Cin); Philadelphia (Phl)).

### Environmental predictors of urban-rural clines in HCN frequencies

On average, urbanization was associated with reduced HCN frequencies across cities, whereby the main effect of standardized distance from the urban center was positively associated with the frequency of HCN within *T. repens* populations (main effect of Distance, F_1,689_ = 36.42, *P* < 0.001, Fig. 2). In a model with unstandardized distance as a predictor, this translated into an average increase in HCN frequency of 0.3 % per km from the urban center. However, the strength of urban-rural phenotypic clines in HCN varied across cities (Distance × City interaction: F_15,689_ = 3.26, *P <* 0.001, Table 1, Fig. 2). The strength of urban-rural clines decreased with increasing values along PC1_Slope_ (R^2^ = 28.4%, *β* = −0.036, *t*_13_ = −2.27, *P* = 0.04, Table 2, Fig. 4), implying that the strongest clines occurred in the warmest environments and the weakest clines occurred in regions of low temperature and high snowfall/depth.

**Figure 4:**
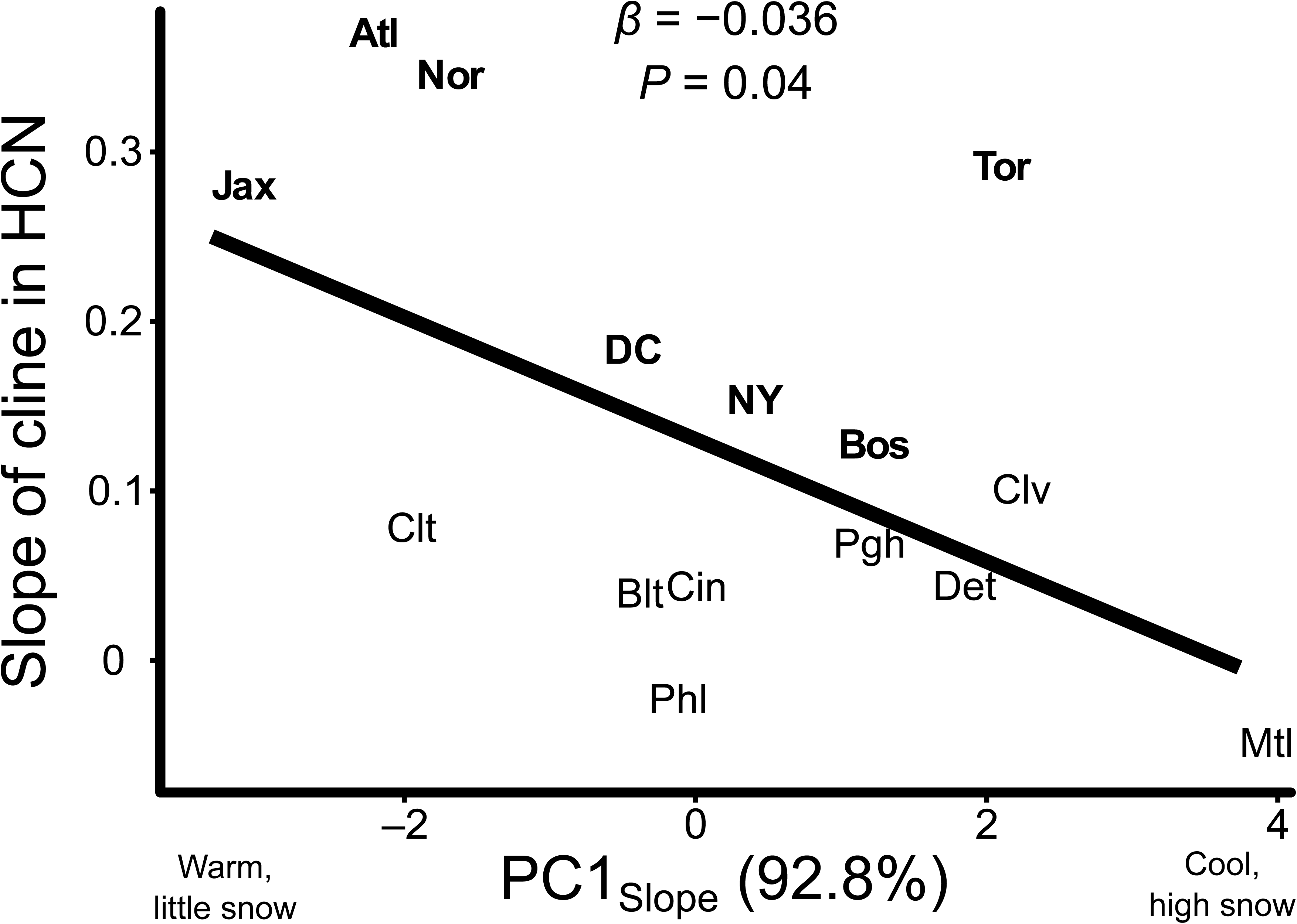
The strength of urban-rural clines in HCN was influenced by PC1_Slope_, a composite axis that accounts for 92.8% of the variation in minimum winter temperature (°C, bio6), maximum summer temperature (°C, bio5), snowfall (cm), and snow depth (cm). City labels are slightly jittered to avoid overlap, if necessary. Bolded cities shower significant linear changes in HCN along urbanization gradients. Cities with low values along PC1_Slope_ have little snow and higher minimum winter and maximum summer temperatures, whereas cities with high values along PC1_Slope_ have the opposite. City abbreviations are the same as in Fig. 3.

### Clines at loci underlying HCN and deletion haplotype frequencies

Of the 16 cities surveyed in this study, we assayed the genotype at the two underlying genes in nine cities, six of which showed significant clines in HCN (Table 1). Of the six cities with significant clines, three (Atlanta, Jacksonville, Toronto) showed significant linear clines at both *Ac* and *Li*, three (New York, Norfolk, Washington) showed significant linear clines only at *Ac*, whereas no cities had a significant linear cline only at *Li*. Significant clines at *Ac* and *Li* were always in the same direction as clines in HCN (i.e., lower frequencies of the dominant alleles *Ac* and *Li* in urban populations). None of the three cities that lacked a cline in HCN showed a cline at either underlying gene.

All deletion haplotypes at *Ac* and *Li* identified previously in this system were found in each city, and their relative frequencies did not vary for either locus between urban and rural populations (fig. S3 and S4). Haplotype diversity (*Ac*: Simpson’s D_rural_ = 0.41, D_urban_ = 0.38, *t*_11_ = −0.23, *P* = 0.82; *Li*: Simpson’s D_rural_ = 0.60, D_urban_ = 0.52, *t*_11_ = −0.74, *P* = 0.47) did not vary across urban and rural habitats for either locus. Together, these results suggest that no specific deletion haplotype is favored in urban habitats at either *Ac* or *Li*.

## Discussion

We combined field sampling of white clover populations from large eastern North American cities with environmental data to assess the environmental predictors of cyanogenesis on a continental scale and of urban-rural gradients in HCN frequencies. Several key results are most important to answering our research questions. As expected, HCN frequencies decreased northward across the continent as frost exposure increased (question 1). Urban-rural cyanogenesis clines occurred in half of the cities studied (question 2) and the strongest clines occurred in the warmest environments (question 3). Clines in HCN were matched by clines at one or both of the loci underlying HCN, and these clines were always in the same direction (question 4). Finally, the diversity of deletion haplotypes among acyanogenic plants was consistent across urban and rural populations of multiple cities (question 5). Together, these results provide compelling evidence that selection is driving parallel evolution of cyanogenesis clines across multiple large urban centers in North America, although regionally cold and snowy climates dampen parallel responses of HCN to urbanization. Below, we discuss our results in the context of the environmental drivers of HCN evolution at the scale of entire continents and individual cities.

### Environmental predictors of HCN frequencies

The cyanogenesis polymorphism has long served as a model for assessing the climatic drivers of adaptation in natural populations. Pioneering work in the 1950’s and 1960’s identified cold winter temperatures as key drivers of reduced HCN frequencies at northern latitudes and higher altitudes (Daday 1954a,b, 1958, 1965). More recent work corroborated the finding that colder environments have reduced cyanogenesis (Ganders 1990; Kooyers and Olsen 2012, 2013) and additionally identified aridity as a correlate of HCN frequencies, with more HCN in drier habitats due to selection favoring plants producing cyanogenic glucosides (*Ac*) (Kooyers and Olsen 2013; Kooyers et al. 2014). Our results are consistent with previous work demonstrating a cost to producing HCN or its metabolic components in frost-prone habitats (Daday 1954a, 1958, 1965; Ganders 1990; Kooyers and Olsen 2013; Kooyers et al. 2018): northern cities with lower winter temperatures and greater frost exposure had reduced cyanogenesis than southern cities. In contrast to previous work (Kooyers and Olsen 2013), we did not identify aridity as an important predictor of mean HCN frequencies, possibly because the latitudinal transect sampled here spanned a shallow aridity gradient (annual aridity index range: 0.84 – 1.22) and steeper gradients may be necessary to detect aridity as an important correlate of HCN frequencies (e.g. New Zealand cline, aridity index range: 0.5941– 4.8569, Kooyers and Olsen 2013).

### Urban-rural clines in cyanogenesis

Although the repeated appearance of clines in different cities suggests that selection is acting on HCN, population genetic simulations demonstrated that genetic drift can also generate similar patterns (Santangelo et al. 2018a). Thus, the presence of repeated clines in HCN is insufficient evidence for the role of selection to drive adaptation of urban populations. Two current lines of evidence suggest a negligible role of genetic drift in driving urban HCN clines. First, observed clines in HCN are substantially stronger than those expected under realistic gradients in the strength of drift alone (Santangelo et al. 2018a), suggesting that other processes are additionally contributing to the presence of clines. Second, recent population genetic analyses show no increased strength of genetic drift in urban white clover populations, and clines are evolving despite substantial gene flow between urban and nonurban populations, consistent with natural selection driving local adaptation of urban populations (Johnson et al. 2018).

Two additional lines of evidence presented here solidify the role of selection rather than drift in driving the formation urban-rural HCN clines. First, the presence of repeated clines in the same direction at individual loci underlying HCN strongly implicates selection since genetic drift is expected to drive random fluctuations in allele frequencies at a single locus (Santangelo et al. 2018a). Second, the negative relationship between the strength of clines and regional winter conditions suggests that latitudinal variation in winter temperature and snow depth—or something correlated with it—modulates the strength of selection along urban-rural gradients, driving phenotypic clines in cyanogenesis. Additionally, the presence of clines at each individual locus, and equivalent deletion haplotype diversity in urban and rural populations across multiple cities, both suggest that selection favors acyanogenic plants rather than alternative biological functions of individual loci or because of genes tightly linked to particular deletion haplotypes.

Environmental heterogeneity among ‘replicate’ environments can reduce the extent of parallel evolution (Stuart et al. 2017). Previous work in white clover identified colder winter temperatures in urban populations as a putative mechanism driving reduced HCN frequencies in urban environments, which was alleviated in cities with high snowfall (Thompson et al. 2016). Based on this earlier work, we predicted the weakest clines in cities with high mean winter temperature or high snowfall due to the absence of frost-mediated selection against HCN in these cities. Consistent with this prediction, cities with high snowfall (i.e. high values along PC1_slope_) had the weakest clines, potentially due to snow buffering plants from frigid temperatures and weakening frost-mediated selection against HCN-producing genotypes. However, the strongest clines occurred in the warmest cities (i.e. low values along PC1_slope_), contrary to our predictions. Importantly, this provides no information about whether lower winter temperatures in cities is an important selective agent; urban frost may still be important in frost-prone cities with shallower urban-rural gradients in snow depth, as suggested by currently available data (Thompson et al. 2016). However, this does suggest that alternative mechanisms must drive the evolution of clines in warmer cities where frost is uncommon. Indeed Atlanta, which gets little snow (mean snowfall = 0.07 cm/year) and is relatively warm throughout the winter months (average minimum winter temperature = −0.43 °C) contained the strongest phenotypic cline in HCN observed in any city to date (*β* = 0.36, Table 1).

The stronger clines in warmer cities suggests that regional temperature modulates the strength of selection along urban-rural clines in some cities. Given that cyanogenesis functions as an antiherbivore defense (Dirzo and Harper 1982; Thompson and Johnson 2016; Santangelo et al. 2018b), some clines may be explained by differential herbivory among urban and rural populations if urbanization reduces herbivore damage (Raupp et al. 2010; Moreira et al. 2019). Although previous experimental work found a negligible role of herbivory as a driver of urban clines in HCN (Thompson et al. 2016), this work was performed in a single northern city (Toronto). Since herbivory often increases with warmer temperatures (Lemoine et al. 2014), the role of herbivory in generating urban-rural clines in HCN may be more important in warmer, southern cities. Additional work quantifying the strength of clover-herbivore interactions, and biotic interactions more generally, in these cities is needed. Alternatively, recent experimental data suggests a cost to producing the metabolic components of HCN in stressful environments, especially for cyanogenic glucosides (Kooyers et al. 2018). If environmental stressors are stronger in cities (e.g. frost, salinity, pollution, etc.), costs involved in the production of the metabolic components of HCN may result in selection against these genes and lower frequencies in urban populations. Consistent with this hypothesis, some urban-rural clines in HCN were mirrored only by clines at *Ac*, suggesting selection may be acting on this locus due to its greater metabolic costs (Kooyers et al. 2018).

### Conclusions and future directions

We have demonstrated the repeated evolution of urban-rural cyanogenesis clines across eastern North American cities. A major goal for future work in this system entails distinguishing among the targets of selection across replicate clines (i.e., *HCN* vs. *Ac*. vs. *Li*) and disentangling the numerous ecological (e.g., environmental factors) and evolutionary (e.g., selection, drift) drivers of (non)parallel responses of HCN to urbanization. This work will require quantifying a broad array of environmental factors at a finer-scale (e.g. population-level) in cities spanning all continents. White clover is a natural model for understanding how cities drive parallel evolution on a global scale due to its ubiquity across the globe and ease of sampling and phenotyping. Such work would advance our understanding of how cities influence the evolution of populations in our own backyards, and further cement the utility of cities as useful models for understanding the causes and consequences of parallel evolution in nature.

## Data accessibility

All data and code will be made publicly available on the Dryad digital repository upon publication. All code and data can additionally be found on the GitHub page for JSS (https://github.com/James-S-Santangelo/uac).

## Acknowledgment

We thank Daniel Anstett, Spencer Barrett, Connor Fitzpatrick, Molly Hetherington-Rauth, Ruth Rivkin, and John Stinchcombe, whose comments and discussions greatly improved this project. JSS was funded by an NSERC PGS-D. This work was additionally funded by NSERC grants to MTJJ and RWN and a Canada Research Chair (CRC) to MTJJ. The authors declare no conflicts of interest.

## Author contributions

JSS conceived of the study with input from KAT and MTJJ. JSS, KAT, and MTJJ collected samples and JSS, BC, and JS collected data. JSS analyzed the data with input from MTJJ and RWN. JSS wrote the first manuscript draft with subsequent feedback from all authors.

