## supplementary text for "Predicting the strength of urban-rural clines in a Mendelian polymorphism along a latitudinal gradient"

**Online supplementary for: Predicting the strength of urban-rural clines in a Mendelian polymorphism along a latitudinal gradient**

**Authors:** James S. Santangelo^1, 2, 3*^, Ken A. Thompson^4^, Beata Cohan^1^, Jibran Syed^1^, Rob W. Ness^1, 2, 3^, and Marc T. J. Johnson^1, 2, 3^

**Contents:**

- Supplementary text:
  - Text S1: Detailed statistical analyses
  - Text S2: Power analyses for sampling design
  - Text S3: Assessing the fit of non-linear clines
- Supplementary figures: Figures S1 – S20
- Supplementary tables: Tables S1 – S5

**Supplementary text**

**Text S1: Detailed statistical analyses**

To assess the environmental predictors of HCN across our latitudinal gradient, we first calculated mean HCN frequencies for each region by averaging HCN frequencies across all urban and rural populations. We then fit a linear model with mean HCN frequency as our response variable and our environmental variables as predictors. However, because many environmental variables were highly correlated (Table S2), we reduced the total number of predictors included in our model using a two-step process similar to Kooyers and Olsen (2013). We first eliminated predictors that on their own did not significantly predict variation in HCN frequencies at *P* < 0.1, which resulted in the elimination of one variable (snow depth). We additionally removed mPET since it was highly correlated with, and is functionally the same as, aPET, and is incorporated into our alternative measure of aridity (i.e. SMD). Of the nine remaining environmental variables, we retained predictors as independent variables if they were not strongly correlated (|r| > 0.7) with any other predictor, which resulted in three variables (annual aridity, soil moisture deficit, and the # of days < 0°C with no snow cover) being retained. The remaining six variables were all strongly correlated (most |r| > 0.8), preventing us from being able to reliably interpret their independent effects at predicting HCN frequencies. We used principal components analysis (PCA) to summarize these six variables into a single new composite variable, PC1_HCN_. The PCA was performed on normally standardized environmental variables (i.e., mean of 0 and standard deviation of 1) using the ‘prcomp’ function from the ‘stats’ R package (R Core Team 2018). The first PC explained 90.2% of the variation in these variables (eigenvalue = 4.5). Because none of the other axes explained substantial variation (< 7%), we retained only the first PC axis. We refer to the first PC axis summarizing variation in the environmental variables retained for the analysis of HCN frequencies as PC1_HCN_. Lower values of PC1_HCN_ characterized cities with higher summer temperatures, higher minimum winter temperatures, higher summer precipitation, greater potential evapotranspiration, and lower snowfall. Our final model was as follows: HCN frequency ~ PC1_HCN_ + # days < 0°C with no snow + annual aridity index + soil moisture deficit. We fit this model using the ‘lm’ function in R and considered this the ‘full’ model, to which we performed model selection and averaging.

To examine which environmental factors best predict mean HCN frequencies, we generated reduced models with all pairwise combinations of predictors and ranked these models by AIC_c_ (Johnson and Omland 2004; Symonds and Moussalli 2011). To obtain parameter estimates and *P*-values, we averaged the model coefficients from all models with ΔAIC_c_ < 2 (Burnham and Anderson 2002; Richards 2005). Model selection and averaging was performed using the ‘dredge’ function in the ‘MuMIn’ R package (Bartoń 2016), which presents both ‘full’ and ‘conditional’ model coefficients. ‘Full’ model coefficients (not to be confused with the ‘full’ model generated before model selection) sets predictors to 0 if they are not included in component models whereas ‘conditional’ coefficients ignore component models where the predictor is absent. Thus, ‘full’ model coefficients are more conservative (Bartoń 2016) and we interpreted these to obtain parameter estimates and *P*-values.

Prior to examining the environmental predictors of clines in HCN and its component genes, we first tested whether, on average, urbanization influenced HCN frequencies. We fit a linear regression with the proportion of cyanogenic plants within each population as the response variable, and city, standardized distance to the urban center, and the city × distance interaction as predictors. We used distance to the urban center as a measure of urbanization as this is highly correlated with % impervious surface (R^2^ = 0.64, Johnson et al. 2018) and sufficiently captures variation in HCN frequencies across urban-rural gradients (Thompson et al. 2016; Johnson et al. 2018). Distances between populations were calculated using the haversine formula (Sinnott 1984). Because urban-rural transects varied in length, we standardized distance to be between 0 (urban-most) and 1 (rural-most). We fit this model using the ‘lm’ function in R and obtained *P*-values from type 3 sums-of-squares computed using the ‘Anova’ function in the R package ‘car’ (Fox and Weisberg 2011). In our model, a significant effect of City suggests that mean HCN frequencies vary across the 16 cities. A significant Distance term means that across all cities, HCN frequencies vary in parallel across the urban-rural transect (i.e. parallel clines in HCN frequencies), while a significant City × Distance interaction indicates the strength or direction of clines in HCN varies across cities. The significant effects of City and the City × Distance interaction in our model (see “Results”) justify an examination of the environmental predictors of mean HCN frequencies and variation in the strength of clines across cities, respectively.

To examine the environmental predictors of variation in the strength of clines, we first fit separate linear models to the within-population HCN, *Ac*, or *Li* frequency data for each city. Each model contained the proportion of cyanogenic plants as the response variable and the standardized distance to the urban center as the predictor. Models were fit using the ‘lm’ function in R. To examine clines in the frequency of *Ac* and *Li*, we performed the same procedure as above but used the inferred frequency of *Ac* and *Li* from HWE as response variables. For each model, we extracted the beta coefficient (i.e. slope) describing the change in the frequency of HCN (or *Ac*/*Li*) per unit increase in the standardized distance to the urban center and used this as our estimate of the strength of the clines. Note that cities that showed significant changes in HCN frequency with distance (Table 1 in main text) were also significant following Bonferroni correction of logistic regressions using data from individual plants (i.e., 1 for HCN+, 0 for HCN−). Despite some cities being better fit by quadratic models (see supplementary text S3: “Assessing the fit of non-linear clines”, Table S3), we used the beta coefficient from a first-order linear regression for all cities to ensure that all slope values represented the linear change in the frequency of HCN with standardized distance to the urban center. We did not run a cline model for Tampa due to the absence of variation in HCN frequencies along the urban-rural transect (mean HCN frequency ~ 99%); thus, this analysis contains 15 cities.

We used the same procedure as above to reduce the number of environmental predictors of the strength of urban-rural clines in HCN. Only four of the 10 environmental variables (MST, MWT, snow depth, and snowfall) on their own significantly predicted the strength of clines in HCN and were retained for further analysis. However, because all of these predictors were highly correlated (all |r| > 0.86), we again used PCA to distill these variables into a smaller set of component axes. The first PC explained 92.8% of the variation in these four variables (eigenvalue = 3.7) while the remaining axes explained little variation (all variation explained < 4%). We thus retained only the loadings from this first PC in our model. We refer to the first PC axis summarizing variation in the environmental variables retained for the analysis of the strength of clines as PC1_Slope_. Cities with low values along PC1_slope_ have little snow and higher minimum winter and higher maximum summer temperatures. Since all environmental variables that predicted the strength of clines were incorporated into PC1_slope_, we did not perform model selection or averaging. Our final model included the strength of urban-rural clines in HCN as the response and PC1_slope_ as the sole predictor. The model was fit using the ‘lm’ function in R.

To examine whether urban and rural populations varied in the composition of deletion haplotypes, we first calculated to relative frequency of each haplotype at the *Ac* and *Li* loci for urban and rural populations for each of the seven cities for which haplotype data were available. To examine variation in deletion haplotypes across urban and rural habitats, we qualitatively examined the relative frequencies of deletion haplotypes in urban and rural habitats across the seven cities for which haplotype data were available. We additionally used the raw counts of deletion haplotypes at each locus to calculate Simpson’s diversity index for deletions in urban and rural habitats for each city. We fit Simpson’s diversity index as the response variable in a linear model with habitat type (i.e. urban vs. rural) as the sole predictor such that a significant effect of habitat suggested differences in deletion haplotype diversity among urban and rural habitats. All analyses were performed in R v. 3.5.0 (R Core Team 2018).

**Text S2: Power analyses for sampling design**

Because we intended to sample urban-rural transects from a large number of cities relative to previous work, we examined whether fewer populations or plants per population could be sampled to save resources without compromising our ability to detect clines in HCN if they exist. To do this, we randomly resampled the data collected by Thompson *et al.* (2016) for the city of Boston , we chose to resample Boston as it showed the weakest cline in HCN, providing a conservative estimate of the number of plants and populations necessary to reproduce urban-rural clines. We resampled the data under two sampling schemes: (1) randomly resampling between 10 and 20 plants within populations, with replacement, or (2) randomly resampling between 10 and 50 populations along the transect, with replacement. We performed 1000 iterations under each sampling scheme, each time running a linear model with within-population HCN frequencies as the response variable and distance to urban center as the sole predictor. We visually examined plots of the change in mean *P*-value of the clines and mean standard deviation of the cline’s slope with the goal of minimizing the change in these quantities relative to the samplimg scheme in Thompson *et al.* (2016) (i.e. 20 plants in each of approximately 40 populations).

Increasing the number of plants sampled per population and the number of populations along the transect decreased the mean *P-*value and mean standard deviation of slopes across simulated clines (fig. S1). The mean *P*-value and standard deviation of the slope was more strongly affected by decreases in the number of populations sampled along the transect than by reducing the number of plants within populations (fig. S1a and S1b vs. S1c and S1d); for this reason, we opted to maximize the number of populations along the transect and sampled 40 populations from each city as in Thompson *et al.* (2016). By contrast, we reduced the number of plants sampled per population from 20 (Thompson *et al.* 2016) to 15, since we considered the minor reduction in power from sampling 15 plants to be outweighed by the time and resources saved by having fewer samples to process.

**Text S3: Assessing the fit of non-linear clines**

As described in text S1, we fit a linear model to each city with the mean within-population HCN frequency (or *Ac* and *Li* frequencies, where available) as a response variable and standardized distance to the urban core as a predictor. However, visual inspection of some of the regression plots suggested that some cities may be better fit by quadratic models rather than first-order regressions. Therefore, for each regression in each city, we additionally fit quadratic models to the within-population HCN, *Ac*, or *Li*, frequencies (i.e. Response ~ std_distance + std_distance^2^). We considered the quadratic model a better fit if it was more that 2 AIC points lower than the AIC score for the first-order regression.

While most cities were best fit by first-order regressions for HCN, *Ac*, and *Li*, some cities were better fit by quadratic cline models (Table S3). In particular, both Charlotte and Jacksonville showed non-linear changes in the frequency of HCN with increasing distance from the urban center (Table S3, Figure S8 and S12). While no cities showed non-linear changes in the frequency of *Ac* with increasing distance from the urban center, Atlanta (Figure S5), Charlotte (Figure S8), Jacksonville (Figure S12) and Washington (Figure S20) all varied non-linearly in the predicted frequency of *Li* along our urbanization gradient (Table S3).

**Supplementary tables**

**Table S1**: Minimum and maximum years, and total number of weather observations for the months of January and February across all 16 cities. Weather data were obtained as daily values from a single international airport in each city.

| City | Airport station | min_year | max_year | count |
| --- | --- | --- | --- | --- |
| Atlanta | Hartsfield International | 1980 | 2015 | 1612 |
| Baltimore | Washington International | 1980 | 2015 | 2133 |
| Boston | Logan International | 1980 | 2004 | 1364 |
| Charlotte | Douglas International | 1980 | 2015 | 2133 |
| Cincinnati | Northern Kentucky International | 1980 | 2015 | 2133 |
| Cleveland | Hopkins International | 1980 | 2015 | 2132 |
| Detroit | Detroit Metropolitan | 1980 | 2015 | 2122 |
| Jacksonville | Jacksonville International | 1980 | 2015 | 1538 |
| Montreal | Pierre Elliott Trudeau International | 1980 | 2015 | 2109 |
| NewYork | La Guardia | 1980 | 2015 | 2133 |
| Norfolk | Norfolk International | 1980 | 2015 | 1120 |
| Philadelphia | Philadelphia International | 1980 | 2015 | 2073 |
| Pittsburgh | Pittsburgh International | 1980 | 2015 | 2132 |
| Tampa | Tampa International | 1980 | 2015 | 2133 |
| Toronto | Lester B. Pearson International | 1980 | 2013 | 2006 |
| Washington D.C. | Dc Dulles International | 1980 | 2015 | 2132 |

**Table S2**: Pearson product-moment correlation coefficients (lower triangle) and associated *P*-values (upper triangle) for all pairwise combinations of environmental variables collected for our analyses. Bolded coefficients and *P-*values are significant at *P* < 0.05.

|  | **Lat** | **Long** | **AI** | **mPET** | **aPET** | **mPrecip** | **MWT** | **MST** | **SMD** | **SD** | **SF** | **Days < 0°C no snow** |
| --- | --- | --- | --- | --- | --- | --- | --- | --- | --- | --- | --- | --- |
| **Lat** |  | 0.058 | **0.042** | **< 0.001** | **< 0.001** | **< 0.001** | **< 0.001** | **< 0.001** | **0.035** | **< 0.001** | **< 0.001** | 0.263 |
| **Long** | 0.484 |  | **0.001** | **0.041** | 0.062 | 0.397 | 0.155 | 0.21 | 0.808 | 0.215 | 0.171 | 0.42 |
| **AI** | **0.513** | **0.751** |  | **0.021** | **0.032** | 0.252 | 0.078 | 0.116 | 0.987 | 0.181 | 0.098 | 0.568 |
| **mPET** | **-0.876** | **-0.516** | **-0.57** |  | **< 0.001** | **0.007** | **< 0.001** | **< 0.001** | 0.618 | **< 0.001** | **< 0.001** | 0.891 |
| **aPET** | **-0.986** | -0.477 | **-0.538** | **0.938** |  | **< 0.001** | **< 0.001** | **< 0.001** | 0.097 | **< 0.001** | **< 0.001** | 0.415 |
| **mPrecip** | **-0.882** | -0.227 | -0.304 | **0.644** | **0.837** |  | **< 0.001** | **< 0.001** | **< 0.001** | **0.014** | **0.003** | 0.06 |
| **MWT** | **-0.983** | -0.373 | -0.453 | **0.806** | **0.953** | **0.905** |  | **< 0.001** | **0.012** | **< 0.001** | **< 0.001** | 0.284 |
| **MST** | **-0.938** | -0.331 | **-0.409** | **0.893** | **0.95** | **0.775** | **0.93** |  | 0.147 | **< 0.001** | **< 0.001** | 0.859 |
| **SMD** | **-0.529** | 0.066 | 0.004 | 0.135 | 0.429 | **0.845** | **0.609** | 0.38 |  | 0.45 | 0.222 | **0.007** |
| **SD** | **0.817** | 0.328 | 0.352 | **-0.821** | **-0.821** | **-0.6** | **-0.82** | **-0.888** | -0.203 |  | **< 0.001** | 0.34 |
| **SF** | **0.896** | 0.36 | 0.429 | **-0.824** | **-0.891** | **-0.694** | **-0.886** | **-0.941** | -0.323 | **0.91** |  | 0.88 |
| **Days < 0°C no snow** | 0.298 | 0.217 | 0.154 | 0.037 | -0.219 | -0.48 | -0.286 | -0.048 | **-0.648** | -0.255 | 0.041 |  |

Abbreviations: Latitude (Lat); longitude (Long); annual aridity index (AI); monthly PET (mPET); annual PET (aPet); monthly precipitation (mPrecip); minimum winter temperature (MWT); maximum summer temperature (MST); soil moisture deficit (SMD); snow depth (SD); snowfall (SF).

**Table S3:** Best-fit model for the change in within-population HCN, *Ac*, or *Li* frequencies along our urbanization gradient. “Linear” implies that HCN, *Ac*, or *Li* frequencies changed in a linear fashion along our urbanization gradient, whereas “quadratic” means the change in the frequency of HCN, *Ac*, or *Li* was non-linear along our transect. ‘NS’ represents non-significant changes in HCN, *Ac*, or *Li* frequencies along our urbanization gradient. Grey cells show cities for which data at the individual loci was not collected.

| **City** | **HCN** | ***Ac*** | ***Li*** |
| --- | --- | --- | --- |
| Atlanta | linear | linear | quadratic |
| Baltimore | NS | NS | NS |
| Boston | linear |  |  |
| Charlotte | quadratic | NS | quadratic |
| Cincinnati | NS |  |  |
| Cleveland | NS | NS | NS |
| Detroit | NS |  |  |
| Jacksonville | quadratic | linear | quadratic |
| Montreal | NS |  |  |
| New York | linear | linear | NS |
| Norfolk | linear | linear | NS |
| Philadelphia | NS |  |  |
| Pittsburgh | NS |  |  |
| Tampa | NS |  |  |
| Toronto | linear | linear | linear |
| Washington, D.C. | linear | linear | quadratic |

**Table S4**: Top two models with ΔAIC_c_ < 2 returned from the model selection performed using ‘dredge’ in R. Shown are the *β* coefficients for terms included in the model, the model R^2^, *F*-statistic, degrees-of-freedom (df), log-likelihood (LL), AIC_c_, ΔAIC_c_ and model weight. Grey cells represent terms not included in the specific model shown. These two models were averaged to produce the “conditional” and “full” model averaged coefficients in Tables S5 and S6, respectively.

|  | **Estimate (*β*)** | | | | |  |  |  |  |  |  |  |
| --- | --- | --- | --- | --- | --- | --- | --- | --- | --- | --- | --- | --- |
| **Model** | **Intercept** | **AI** | **# Days < 0°C no snow** | **PC1_HCN_** | **SMD** | **R^2^** | **F** | **df** | **LL** | **AIC_c_** | **ΔAIC_c_** | **weight** |
| 1 | 0.983 | -0.292 | -0.015 | -0.061 |  | 0.957 | 89.279 | 5 | 25.882 | -35.763 | 0.000 | 0.527 |
| 2 | 0.694 |  | -0.015 | -0.068 |  | 0.941 | 104.262 | 4 | 23.372 | -35.107 | 0.656 | 0.379 |

Abbreviations: annual aridity index (AI); soil moisture deficit (SMD)

**Table S5:** “Full” model averaged coefficients from the model selection and averaging of top models with environmental predictors of mean HCN frequencies. Top two models were within 2 AIC_c_ points of one another (Table S4).

| **Term** | ***β*** | **SE** | **𝑧** | ***P*** |
| --- | --- | --- | --- | --- |
| **Annual aridity** | -0.170 | 0.179 | 0.912 | 0.362 |
| **# days < 0 °C no snow** | **-0.015** | **0.002** | **8.774** | **< 0.001** |
| **PC1_HCN_** | **-0.064** | **0.008** | **6.987** | **< 0.001** |

**Supplementary figures**

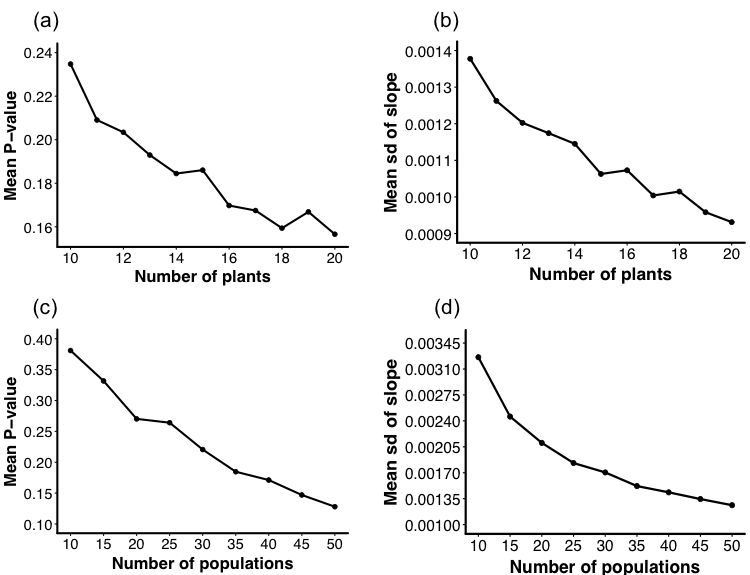

**Figure S1:** Change in mean *P-*value (a and c) and mean standard deviation of the slope of clines (b and d) with increasing number of plants sampled per population (a and b) or increasing number of populations sampled per transect (c and d). Points represent means following 1000 resamplings of plants or populations from the city of Boston (collected by Thompson *et al.* 2016). Slopes and *P-*values are based on regressions using within-population HCN frequency as the response variable and distance to the urban center as the sole predictor.

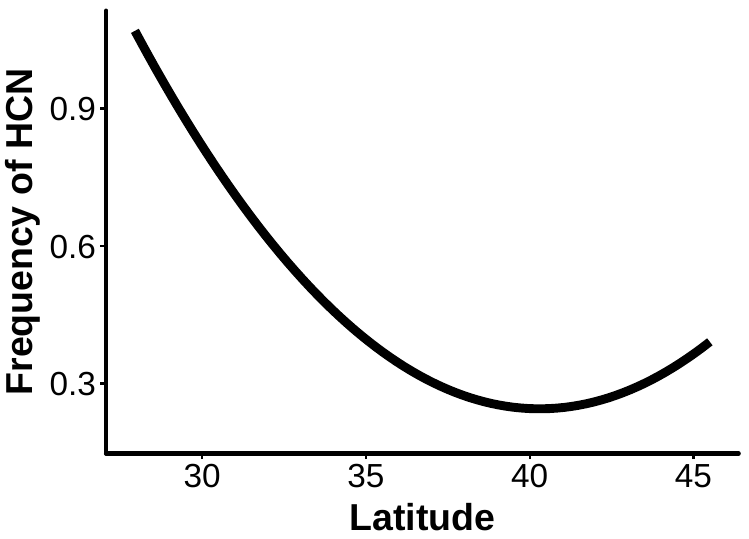

**Figure S2:** Mean frequency of HCN within a city as a function of latitude. HCN frequency is highest in more southern and norther populations. The quadratic model provided a significantly better fit than the linear model based on AIC_c_ (AIC_c_linear_ = −8.8, AIC_c_quadratic_ = −32.8).

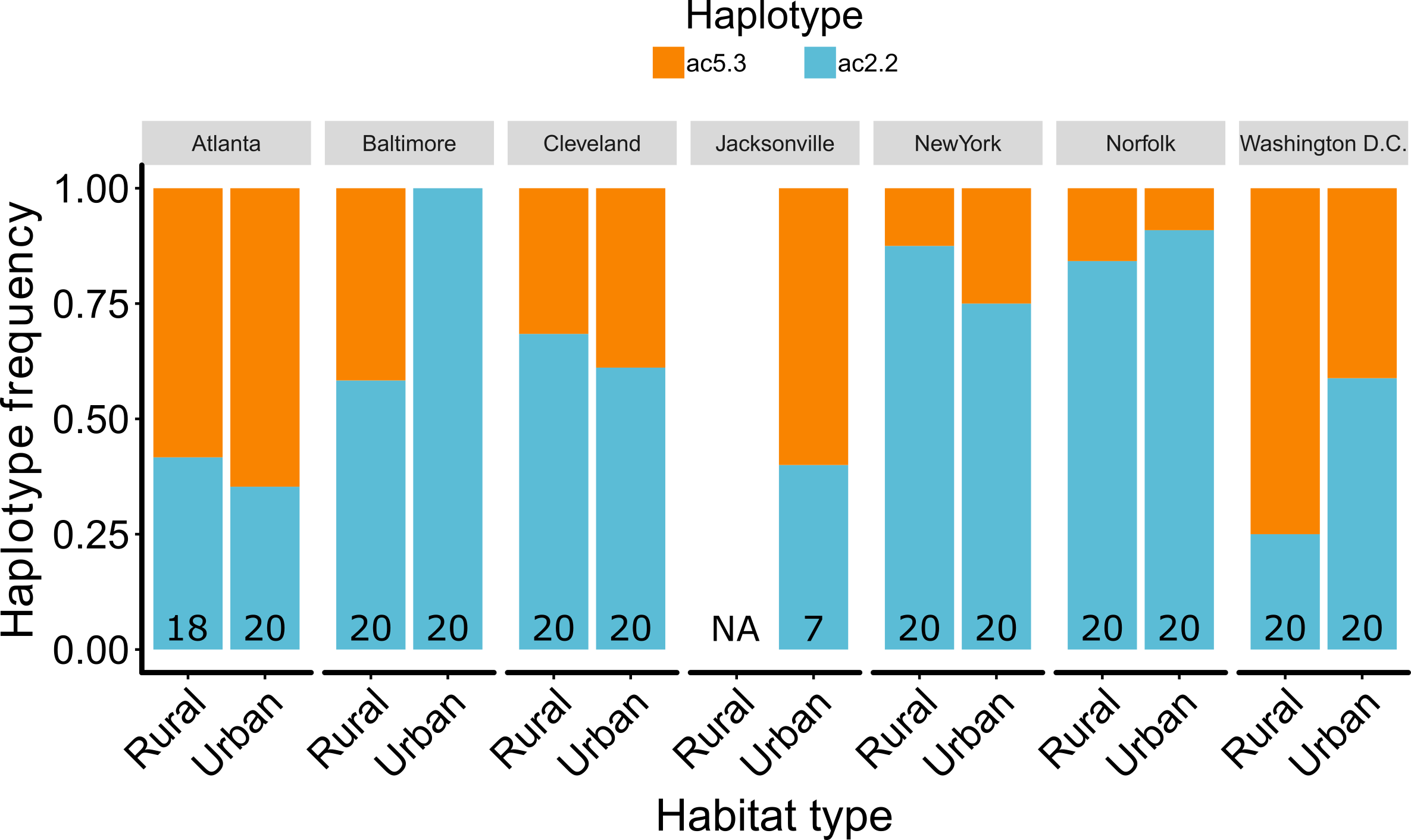

**Figure S3:** Frequency of the 5.3 kbp (ac5.3, orange) and 2.2 kbp (ac2.2, blue) deletion haplotypes at the *Ac* locus (i.e. *CYP79D15*) in urban and rural populations of the seven cities for which haplotype data were sampled. Number of plants assayed for deletion haplotypes are given at the base of bars (note there were no acyanogenic plants in rural Jacksonville that could be assayed). Haplotype naming follows the convention established by Olsen *et al.* (2013) and Kooyers and Olsen (2014).

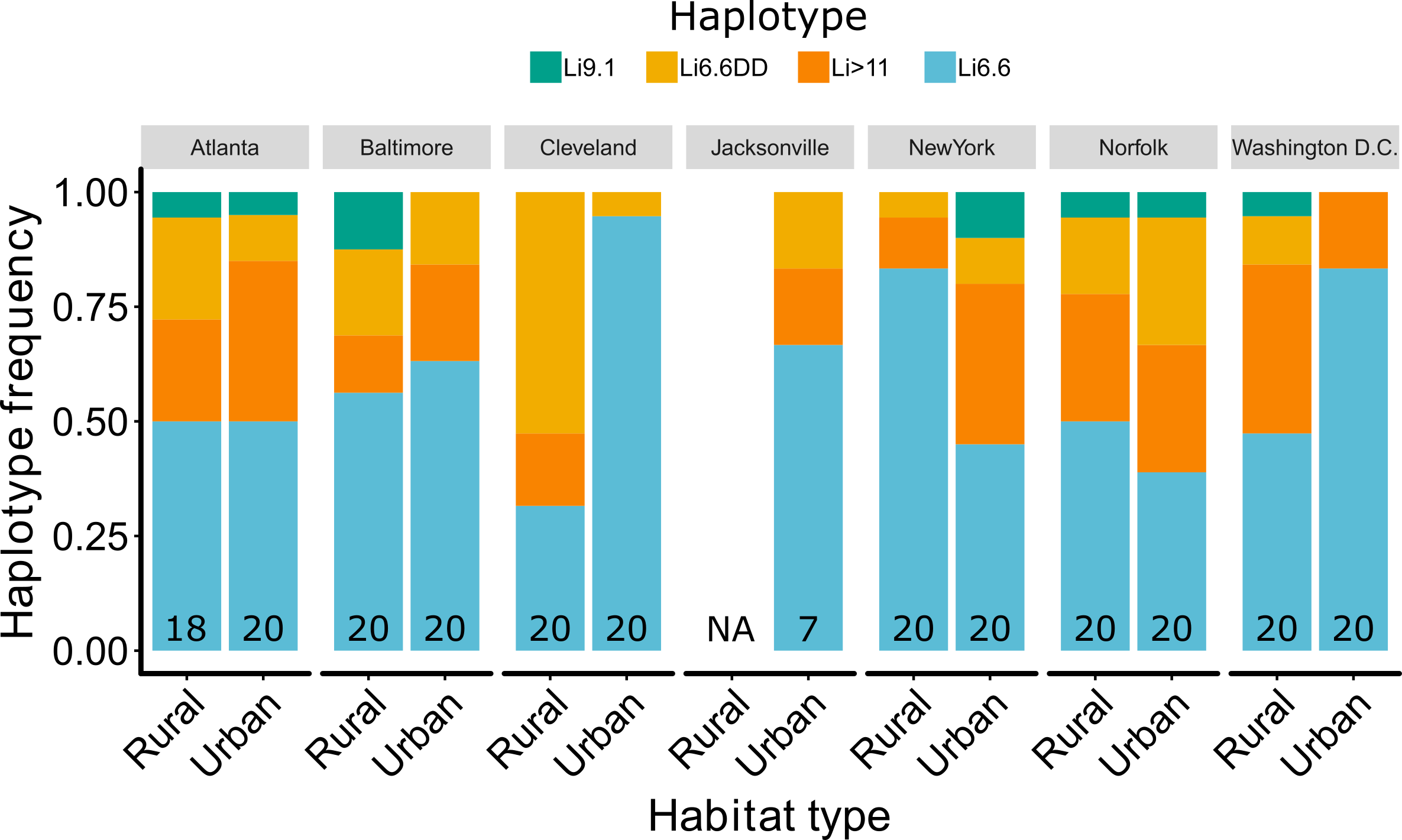

**Figure S4:** Frequency of the 9.1 kbp (Li9.1, green), 6.6 kbp (Li6.6DD, yellow, double deletion), greater than 11 kbp (Li > 11, orange), and 6.6 kbp (Li6.6, blue, single deletion) deletion haplotypes at the *Li* locus in urban and rural populations of the seven cities for which haplotype data were sampled. Number of plants assayed for deletion haplotypes are given at the base of bars (note there were no acyanogenic plants in rural Jacksonville that could be assayed).. Haplotype naming follows the convention established by Olsen *et al.* (2013) and Kooyers and Olsen (2014).

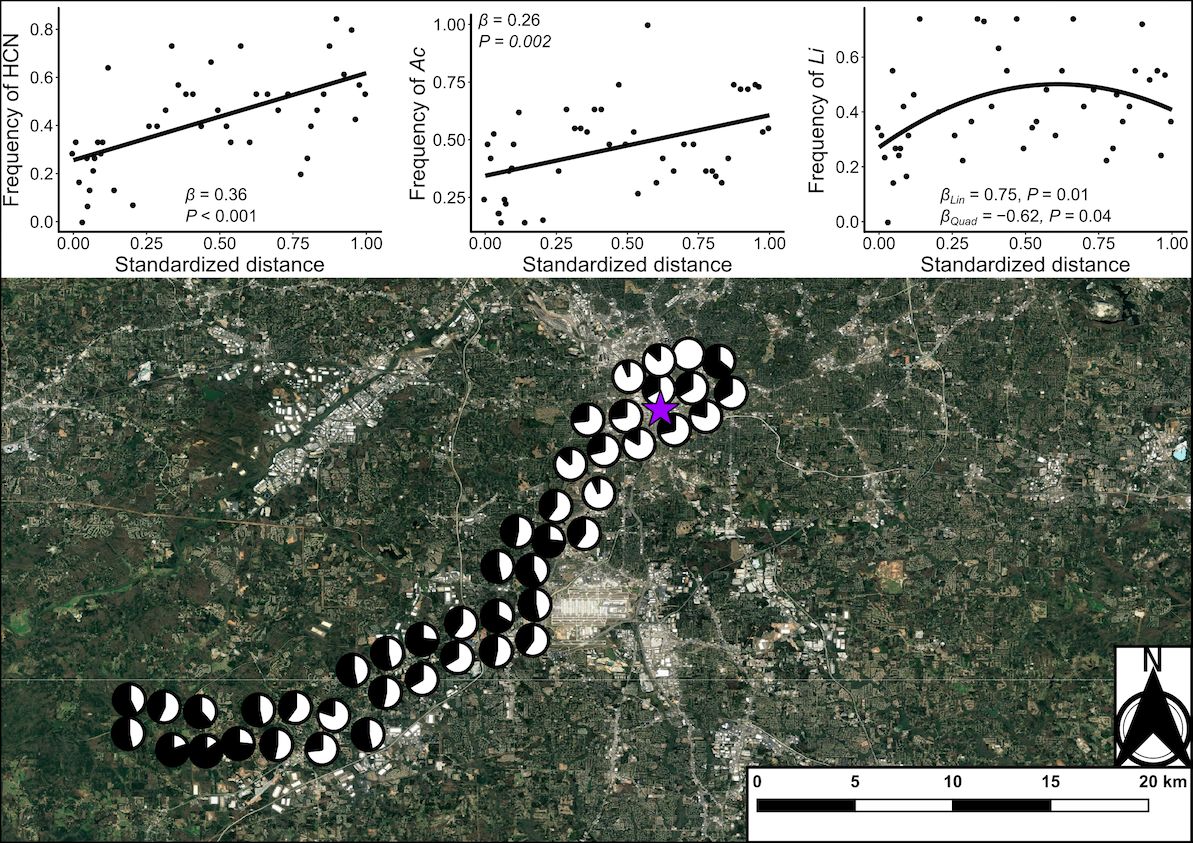

**Figure S5**: Map of the urban-rural transect for the city of Atlanta. Populations along the transect are represented with pie charts showing the proportion of cyanogenic plants (black) in the population. Pie charts have been jittered from their actual location to improve visualization. The purple star represents the location of the city center (Lat: 33.748997, Long: −84.387985). Inset shows the best fit regressions for the change in the frequency of HCN, *Ac*, and *Li* along an urbanization gradient, using standardized distance to the city center as a predictor. For each cline, slopes (*β*) and *P*-values for first-order (linear) and second-order (quadratic, where applicable) terms are provided.

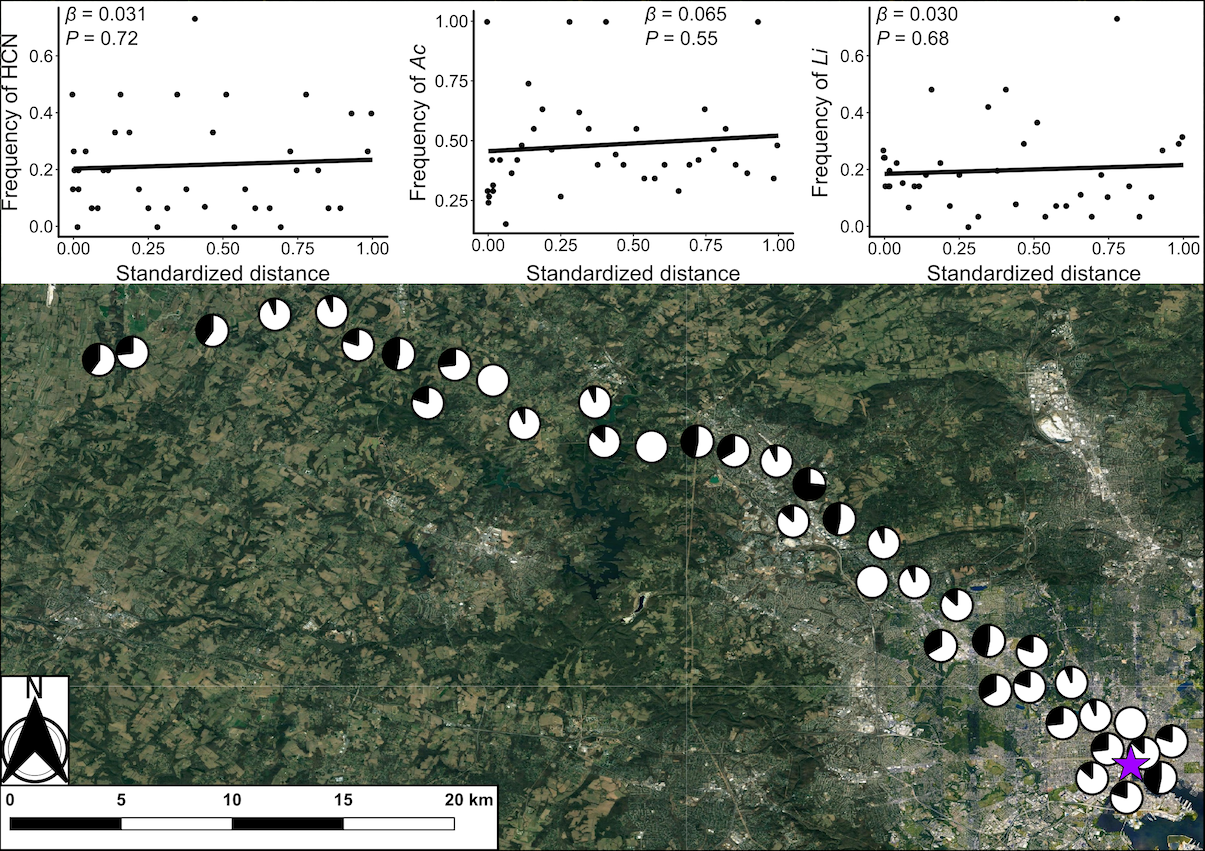

**Figure S6**: Map of the urban-rural transect for the city of Baltimore. Populations along the transect are represented with pie charts showing the proportion of cyanogenic plants (black) in the population. Pie charts have been jittered from their actual location to improve visualization. The purple star represents the location of the city center (Lat: 39.29039, Long: −76.61219). Inset shows the best fit regressions for the change in the frequency of HCN, *Ac*, and *Li* along an urbanization gradient, using standardized distance to the city center as a predictor. For each cline, slopes (*β*) and *P*-values for first-order (linear) terms are provided.

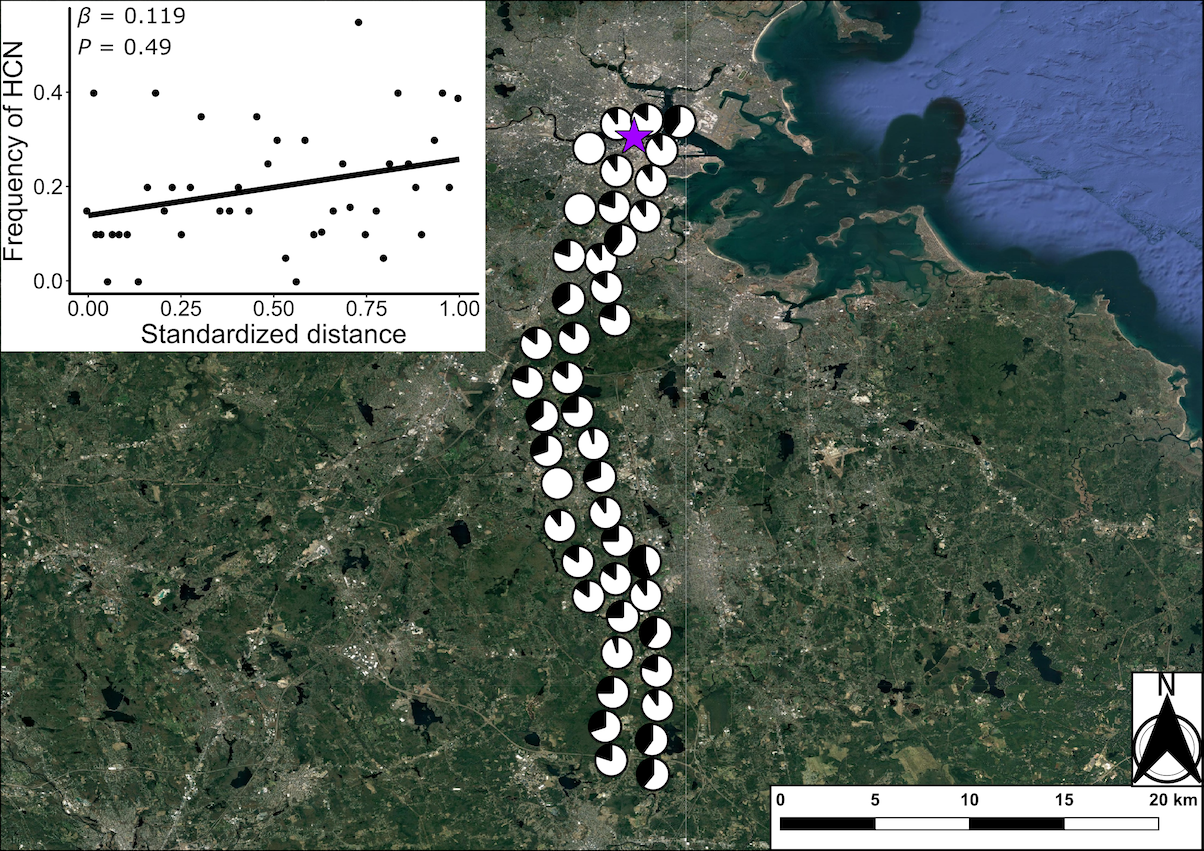

**Figure S7**: Map of the urban-rural transect for the city of Boston. Populations along the transect are represented with pie charts showing the proportion of cyanogenic plants (black) in the population. Pie charts have been jittered from their actual location to improve visualization. The purple star represents the location of the city center (Lat: 42.3547, Long: −71.0665). Inset shows the best fit regressions for the change in the frequency of HCN along an urbanization gradient, using standardized distance to the city center as a predictor. The slope (*β*) and *P*-value for first-order (linear) regression is provided.

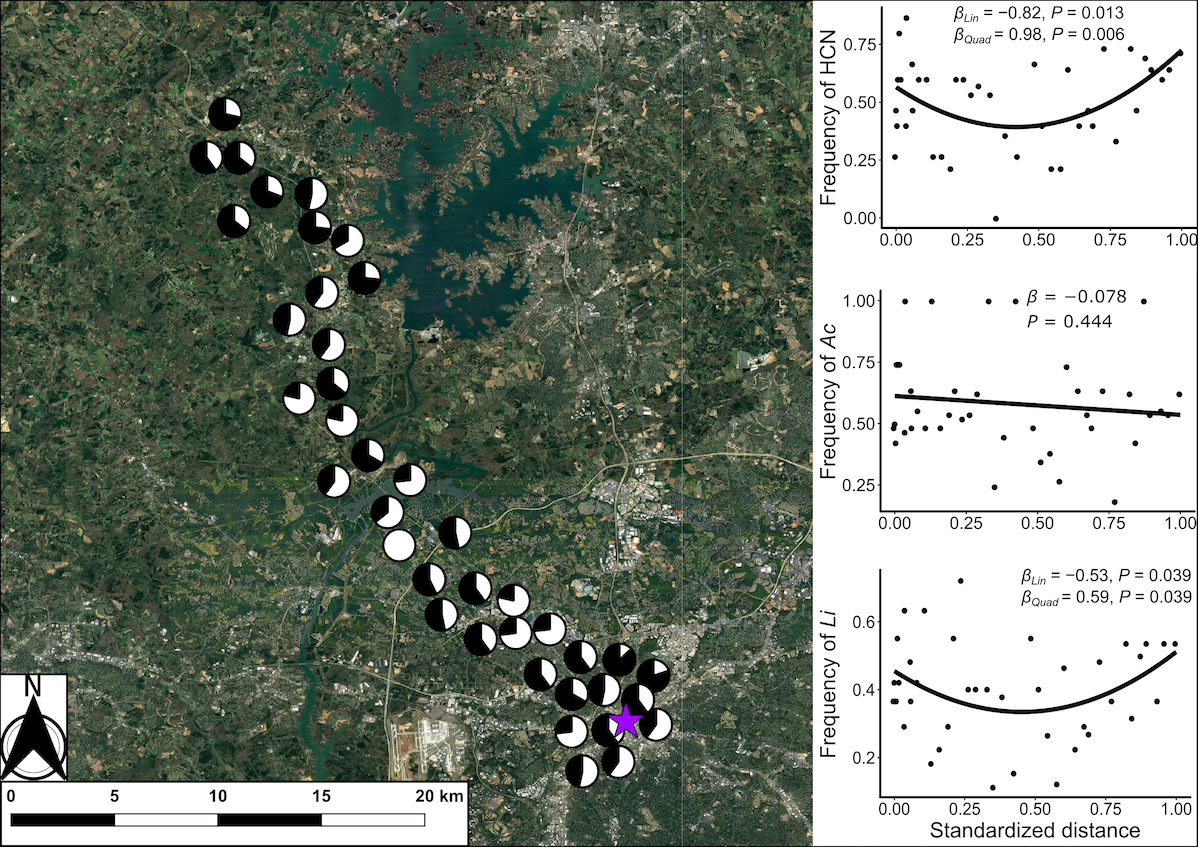

**Figure S8**: Map of the urban-rural transect for the city of Charlotte. Populations along the transect are represented with pie charts showing the proportion of cyanogenic plants (black) in the population. Pie charts have been jittered from their actual location to improve visualization. The purple star represents the location of the city center (Lat: 35.227085, Long: −80.843124). Inset shows the best fit regressions for the change in the frequency of HCN, *Ac*, and *Li* along an urbanization gradient, using standardized distance to the city center as a predictor. For each cline, slopes (*β*) and *P*-values for first-order (linear) and second-order (quadratic, where applicable) terms are provided.

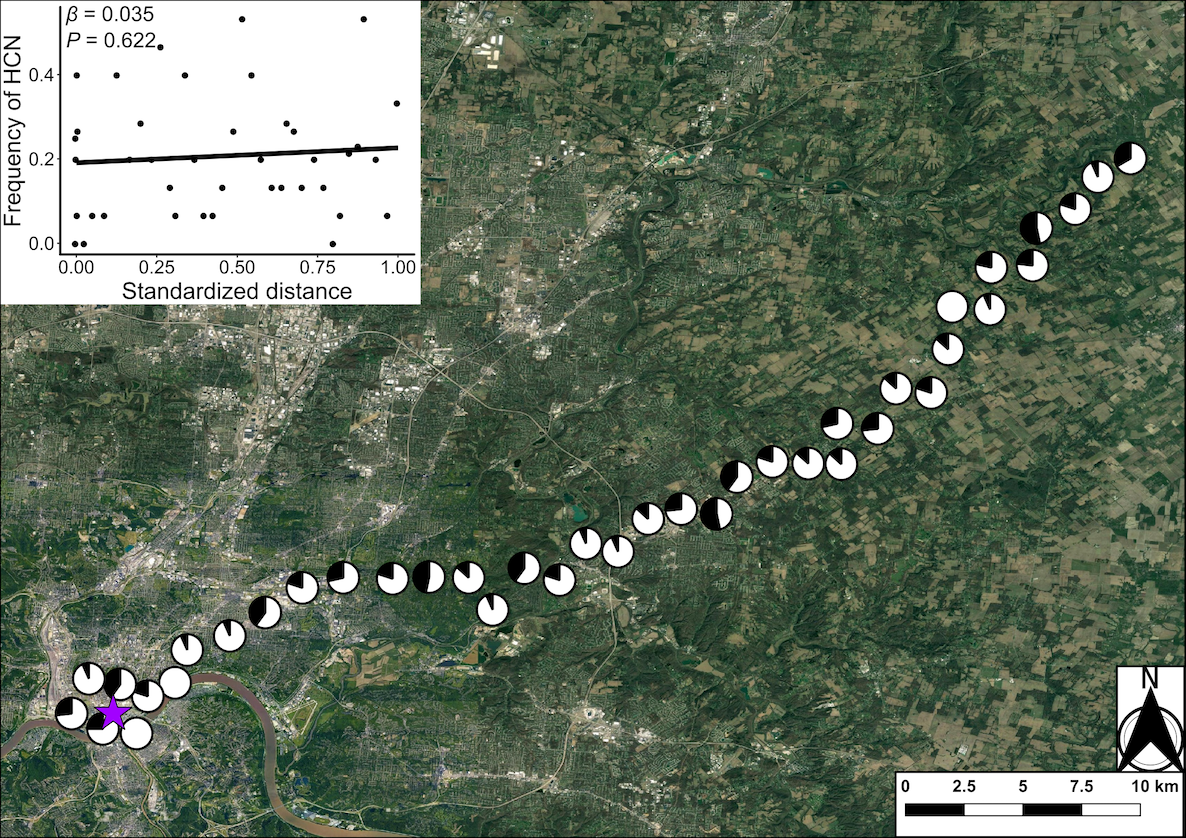

**Figure S9**: Map of the urban-rural transect for the city of Cincinnati. Populations along the transect are represented with pie charts showing the proportion of cyanogenic plants (black) in the population. Pie charts have been jittered from their actual location to improve visualization. The purple star represents the location of the city center (Lat: 39.103119, Long: −84.512016). Inset shows the best fit regressions for the change in the frequency of HCN along an urbanization gradient, using standardized distance to the city center as a predictor. The slope (*β*) and *P*-value for first-order (linear) regression is provided.

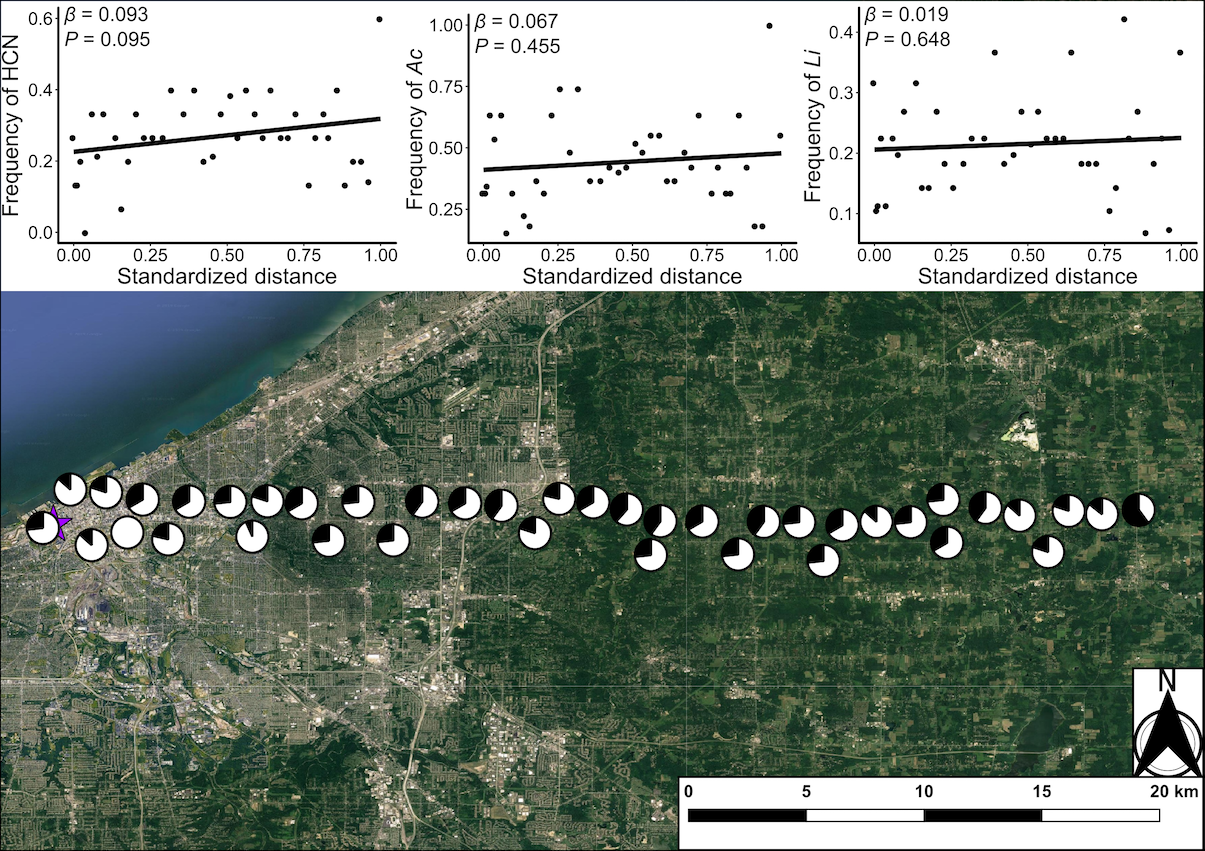

**Figure S10**: Map of the urban-rural transect for the city of Cleveland. Populations along the transect are represented with pie charts showing the proportion of cyanogenic plants (black) in the population. Pie charts have been jittered from their actual location to improve visualization. The purple star represents the location of the city center (Lat: 41.499321, Long: −81.694359). Inset shows the best fit regressions for the change in the frequency of HCN, *Ac*, and *Li* along an urbanization gradient, using standardized distance to the city center as a predictor. For each cline, slopes (*β*) and *P*-values for first-order (linear) terms are provided.

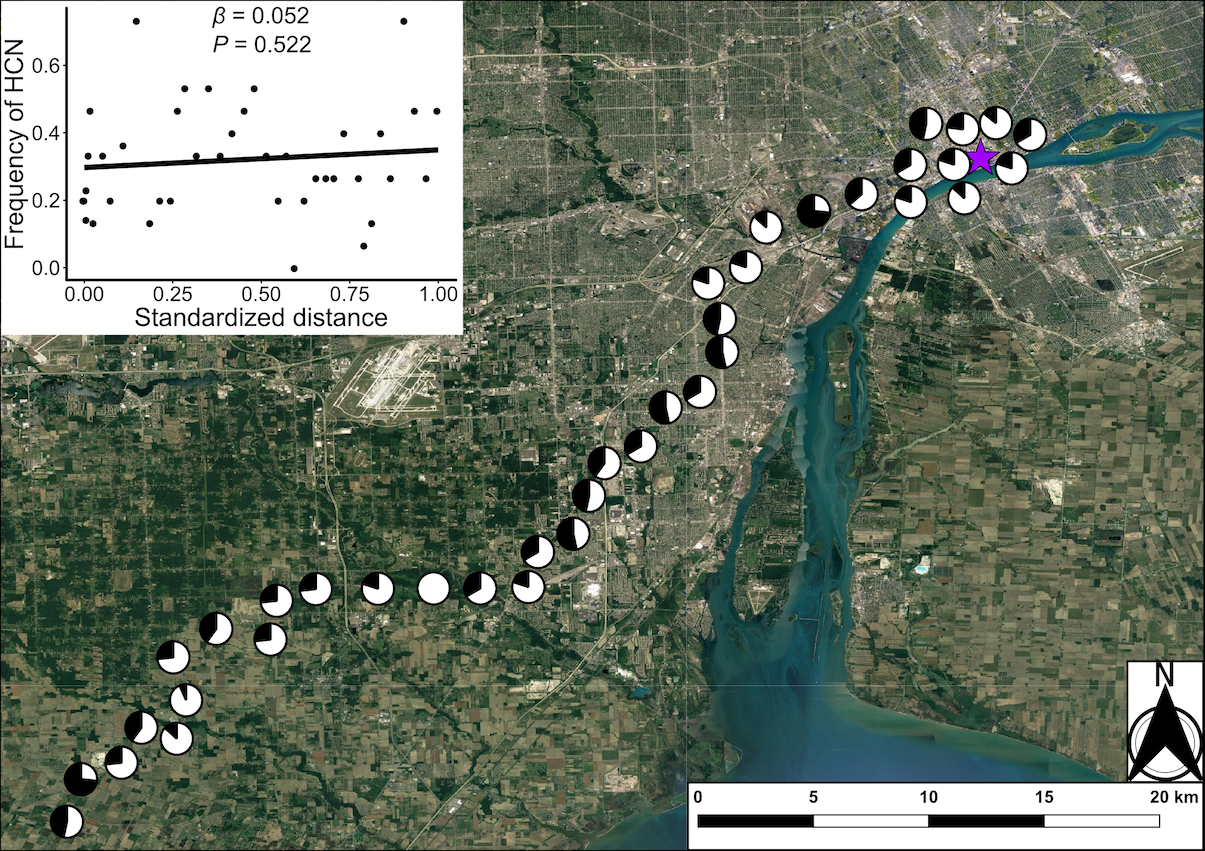

**Figure S11**: Map of the urban-rural transect for the city of Detroit. Populations along the transect are represented with pie charts showing the proportion of cyanogenic plants (black) in the population. Pie charts have been jittered from their actual location to improve visualization. The purple star represents the location of the city center (Lat: 42.331429, Long: −83.045753). Inset shows the best fit regressions for the change in the frequency of HCN along an urbanization gradient, using standardized distance to the city center as a predictor. The slope (*β*) and *P*-value for first-order (linear) regression is provided.

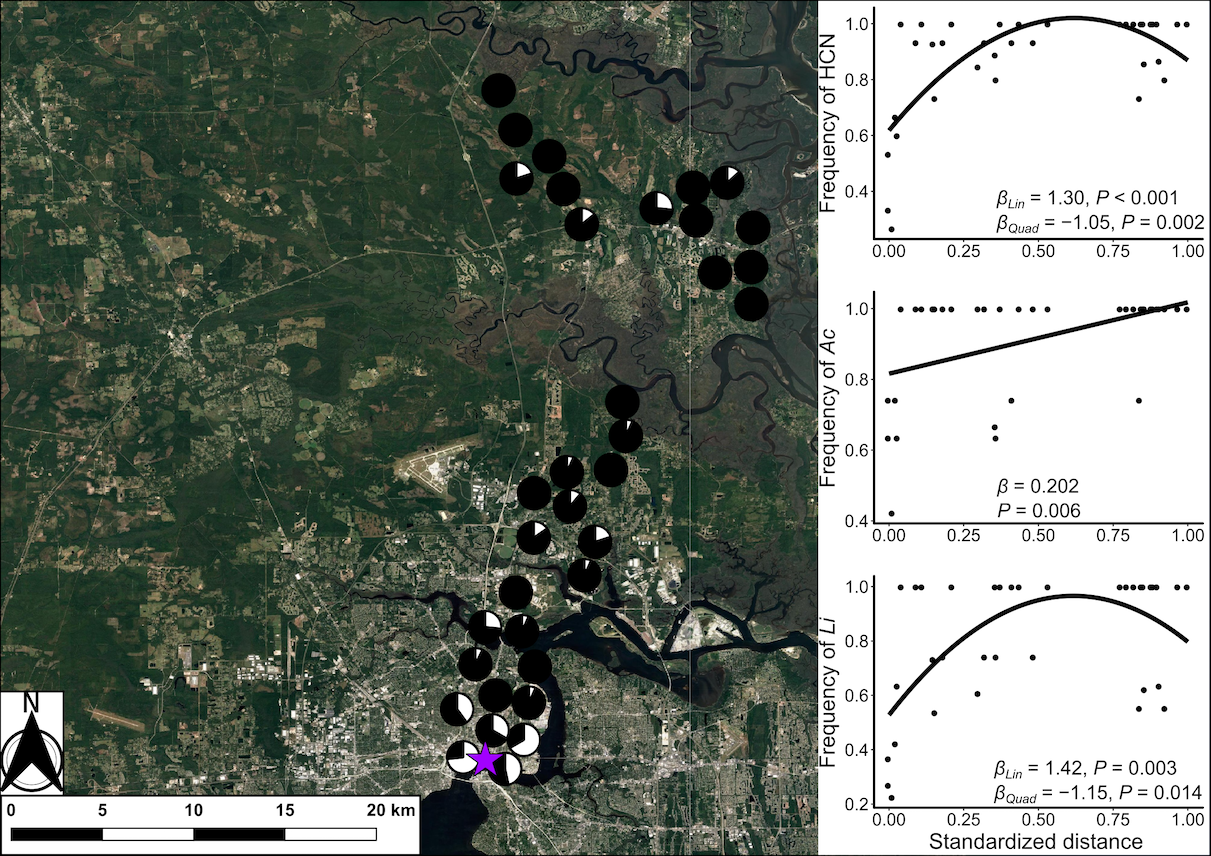

**Figure S12**: Map of the urban-rural transect for the city of Jacksonville. Populations along the transect are represented with pie charts showing the proportion of cyanogenic plants (black) in the population. Pie charts have been jittered from their actual location to improve visualization. The purple star represents the location of the city center (Lat: 30.32597, Long: − 81.656761). Inset shows the best fit regressions for the change in the frequency of HCN, *Ac*, and *Li* along an urbanization gradient, using standardized distance to the city center as a predictor. For each cline, slopes (*β*) and *P*-values for first-order (linear) and second-order (quadratic, where applicable) terms are provided.

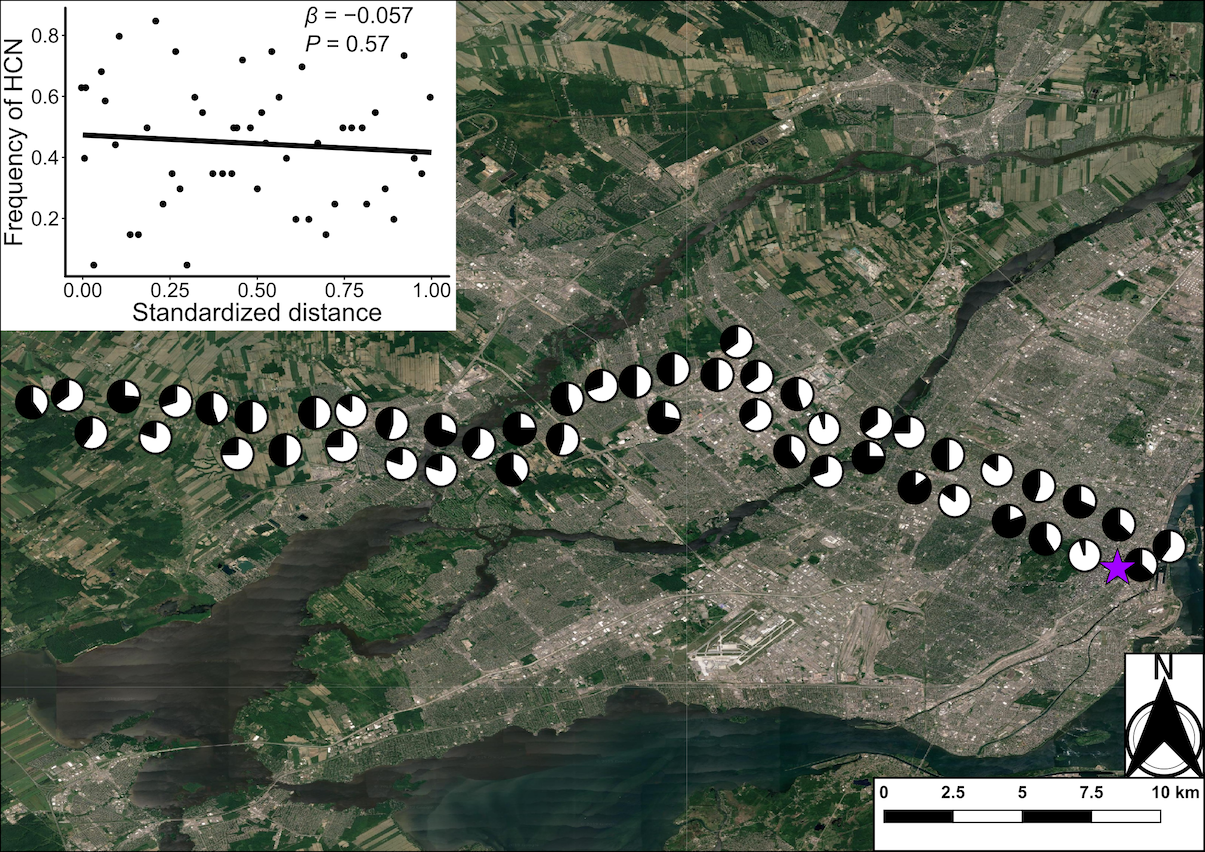

**Figure S13**: Map of the urban-rural transect for the city of Montreal. Populations along the transect are represented with pie charts showing the proportion of cyanogenic plants (black) in the population. Pie charts have been jittered from their actual location to improve visualization. The purple star represents the location of the city center (Lat: 45.502, Long: −73.5672). Inset shows the best fit regressions for the change in the frequency of HCN along an urbanization gradient, using standardized distance to the city center as a predictor. The slope (*β*) and *P*-value for first-order (linear) regression is provided.

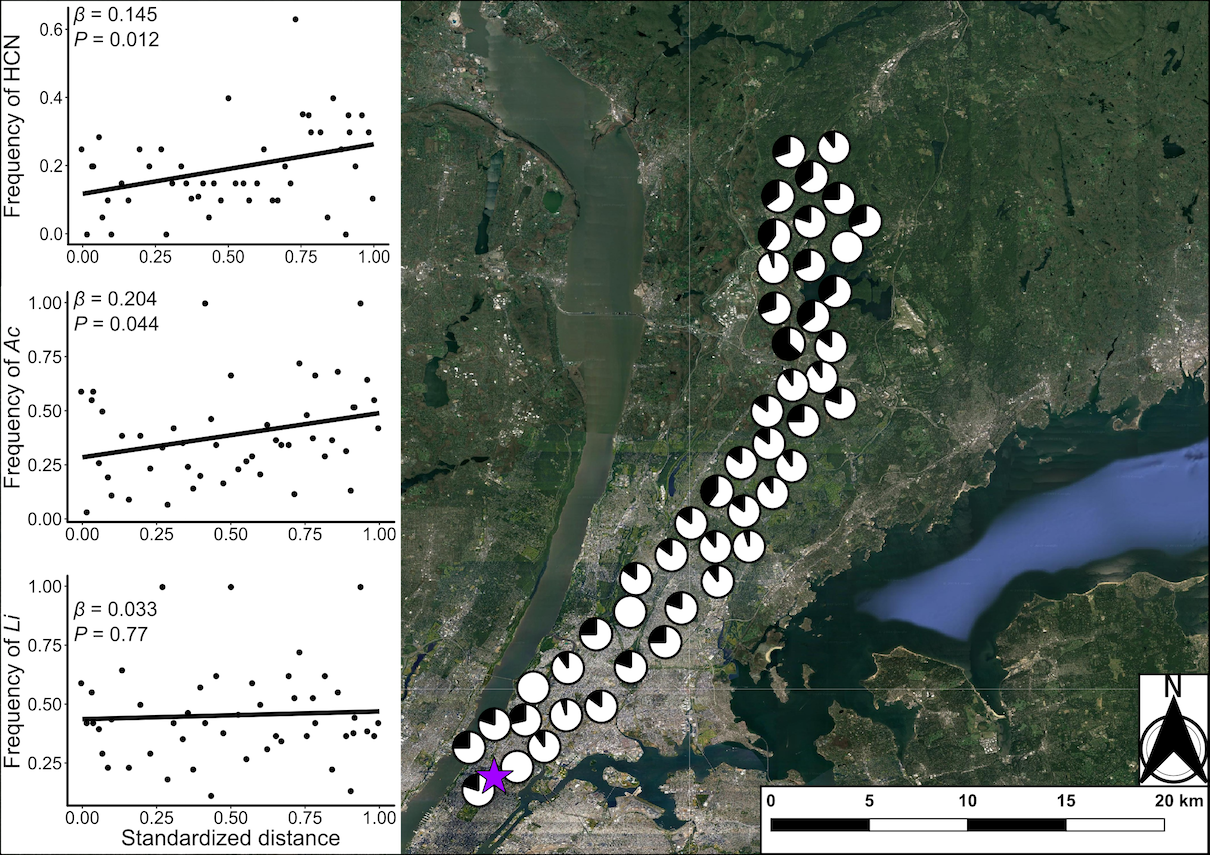

**Figure S14**: Map of the urban-rural transect for the city of New York. Populations along the transect are represented with pie charts showing the proportion of cyanogenic plants (black) in the population. Pie charts have been jittered from their actual location to improve visualization. The purple star represents the location of the city center (Lat: 40.7921, Long: −73.958). Inset shows the best fit regressions for the change in the frequency of HCN, *Ac*, and *Li* along an urbanization gradient, using standardized distance to the city center as a predictor. For each cline, slopes (*β*) and *P*-values for first-order (linear) terms are provided.

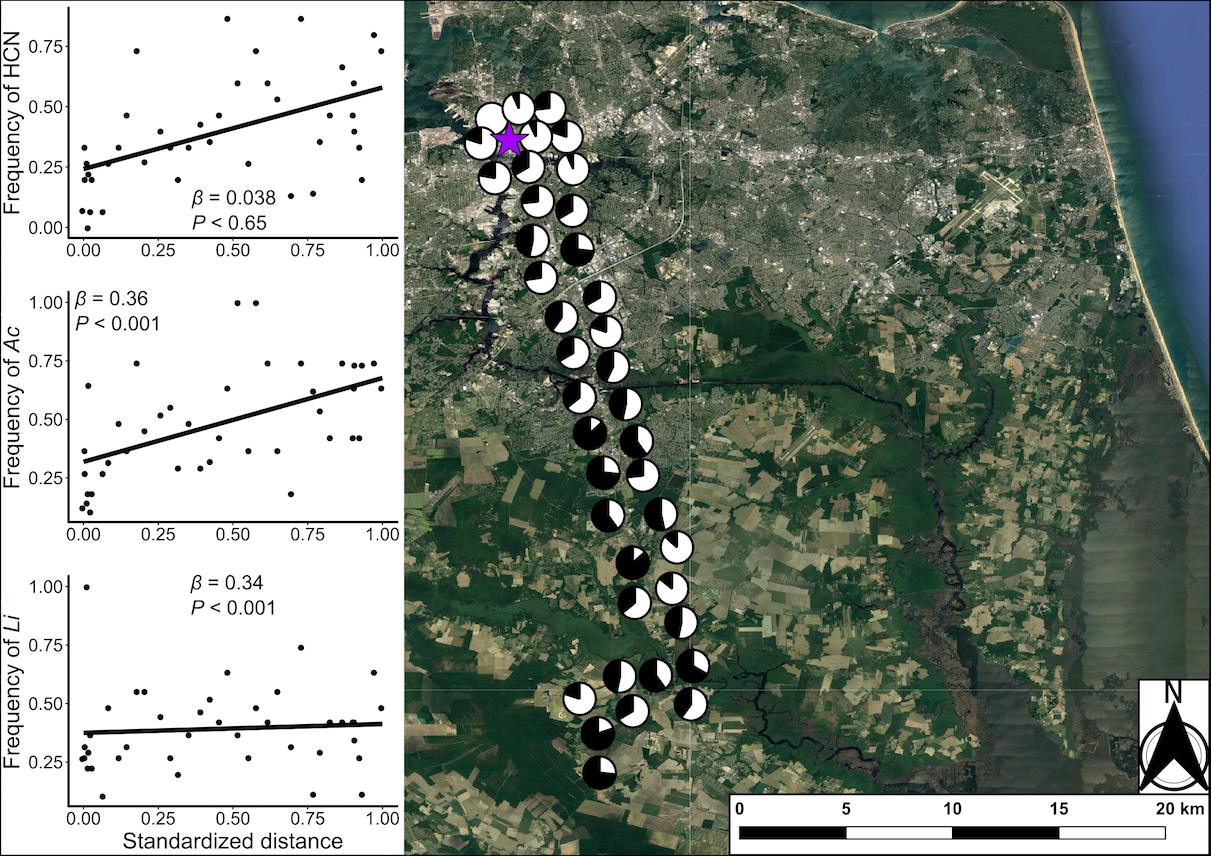

**Figure S15**: Map of the urban-rural transect for the city of Norfolk. Populations along the transect are represented with pie charts showing the proportion of cyanogenic plants (black) in the population. Pie charts have been jittered from their actual location to improve visualization. The purple star represents the location of the city center (Lat: 36.850769, Long: −76.285873). Inset shows the best fit regressions for the change in the frequency of HCN, *Ac*, and *Li* along an urbanization gradient, using standardized distance to the city center as a predictor. For each cline, slopes (*β*) and *P*-values for first-order (linear) terms are provided.

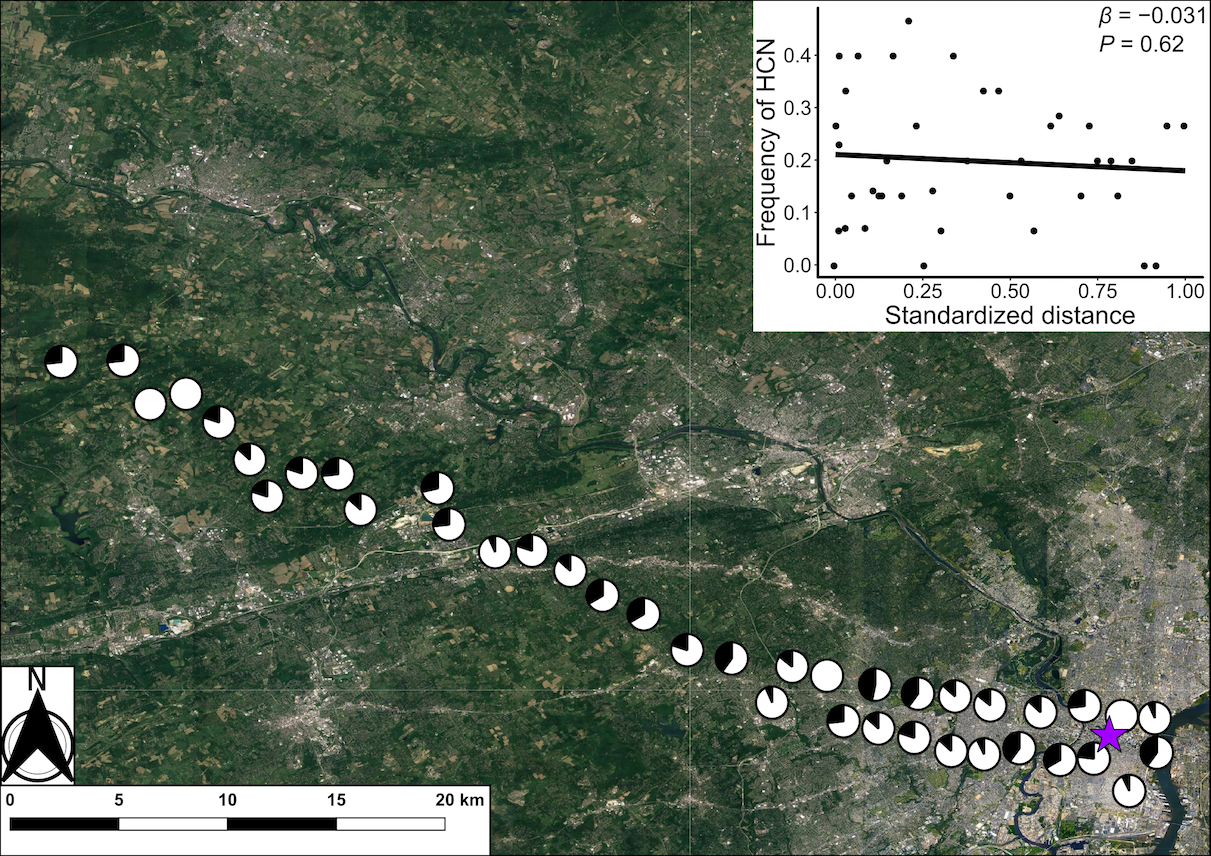

**Figure S16**: Map of the urban-rural transect for the city of Philadelphia. Populations along the transect are represented with pie charts showing the proportion of cyanogenic plants (black) in the population. Pie charts have been jittered from their actual location to improve visualization. The purple star represents the location of the city center (Lat 39.952583, Long: −75.165222). Inset shows the best fit regressions for the change in the frequency of HCN along an urbanization gradient, using standardized distance to the city center as a predictor. The slope (*β*) and *P*-value for first-order (linear) regression is provided.

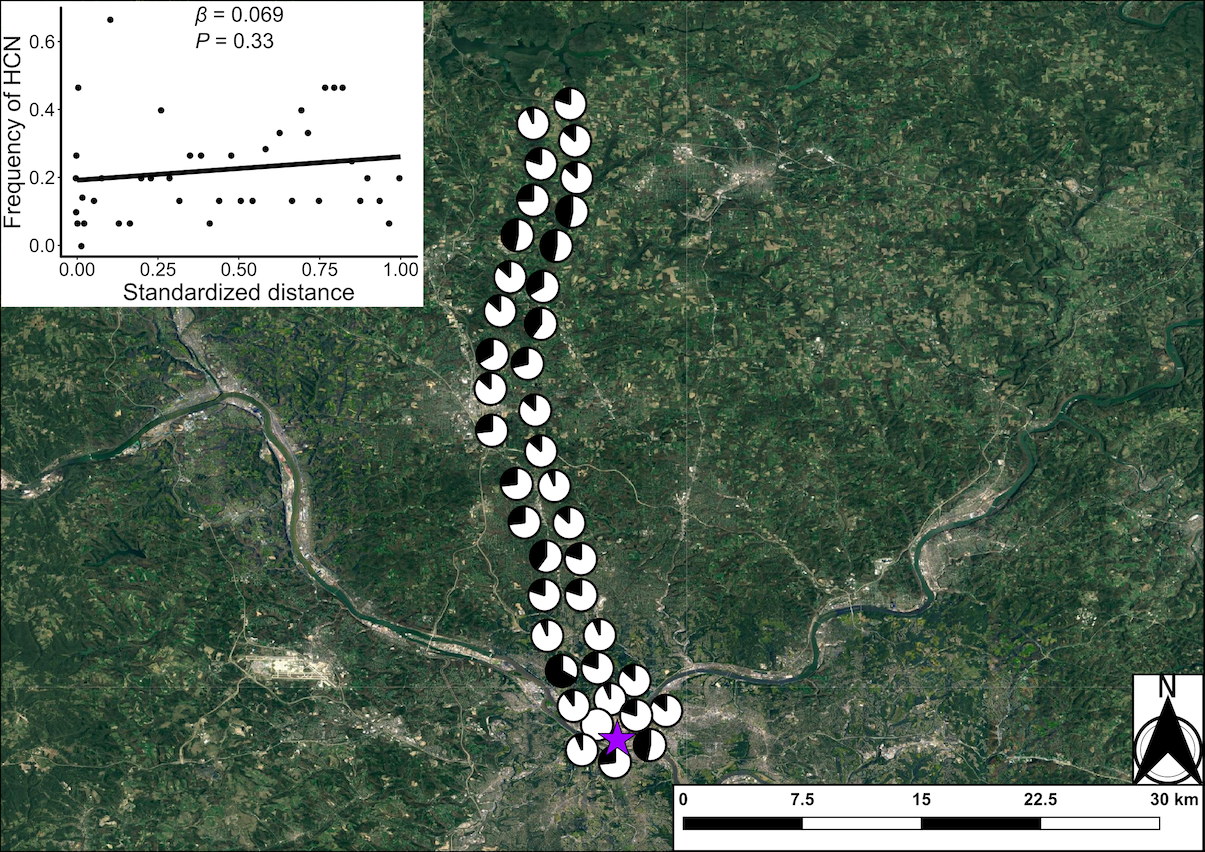

**Figure S17**: Map of the urban-rural transect for the city of Pittsburgh. Populations along the transect are represented with pie charts showing the proportion of cyanogenic plants (black) in the population. Pie charts have been jittered from their actual location to improve visualization. The purple star represents the location of the city center (Lat 40.440624, Long: −79.995888). Inset shows the best fit regressions for the change in the frequency of HCN along an urbanization gradient, using standardized distance to the city center as a predictor. The slope (*β*) and *P*-value for first-order (linear) regression is provided.

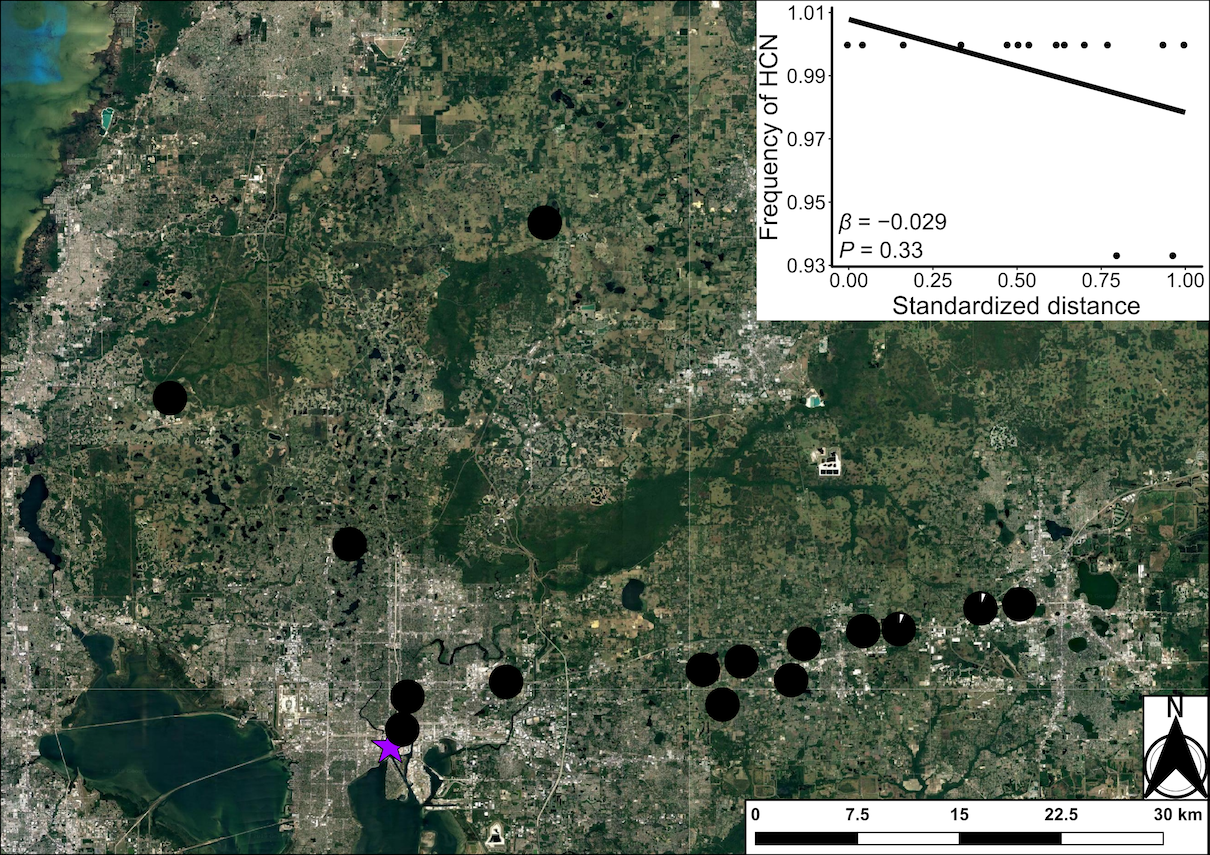

**Figure S18**: Map of the urban-rural transect for the city of Tampa. Populations along the transect are represented with pie charts showing the proportion of cyanogenic plants (black) in the population. Pie charts have been jittered from their actual location to improve visualization. The purple star represents the location of the city center (Lat 27.94742, Long: −82.458778). Inset shows the best fit regressions for the change in the frequency of HCN along an urbanization gradient, using standardized distance to the city center as a predictor. The slope (*β*) and *P*-value for first-order (linear) regression is provided. Note that Tampa was not included in our analysis predicting the strength of clines due to being functionally fixed for cyanogenesis.

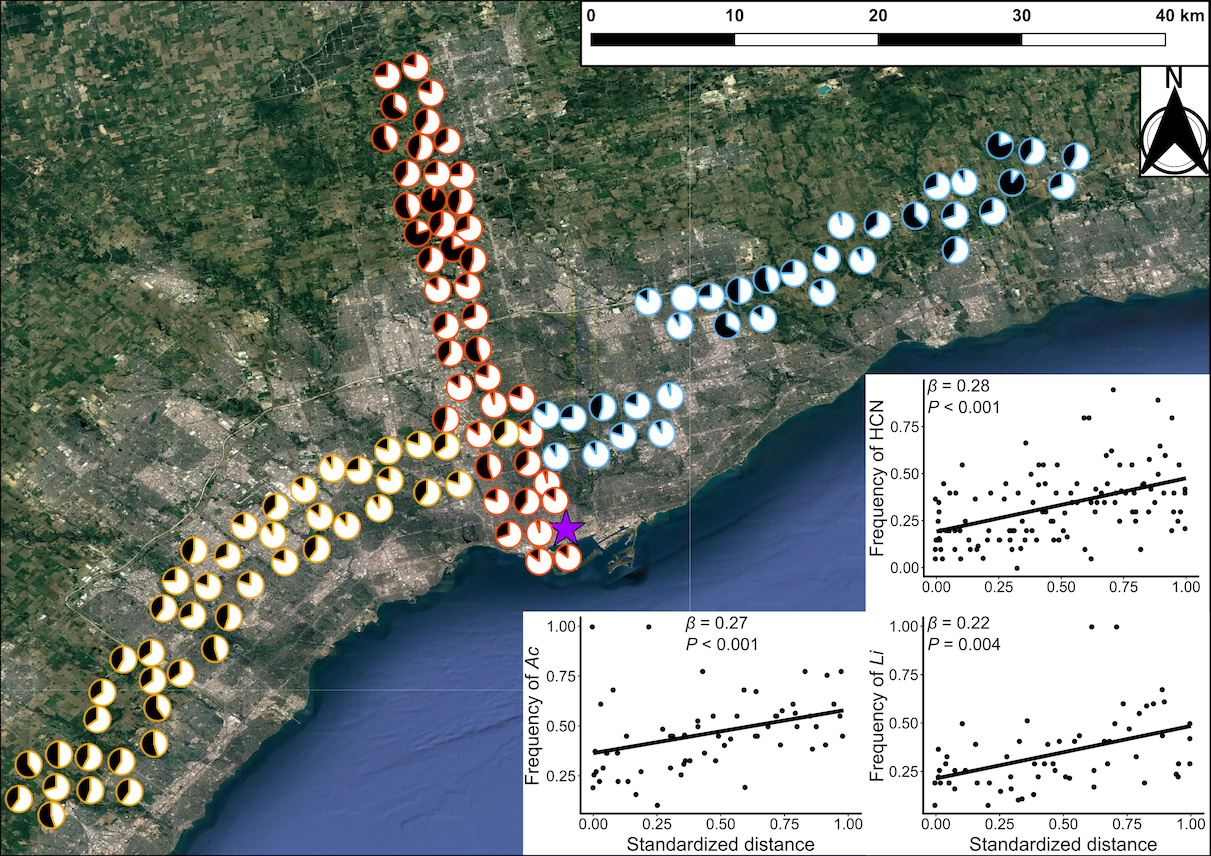

**Figure S19**: Map of the urban-rural transect for the city of Toronto. Populations along the transect are represented with pie charts showing the proportion of cyanogenic plants (black) in the population. Pie charts have been jittered from their actual location to improve visualization. Coloured outlines around pie charts represent the western (yellow), northern (orange), and eastern (blue) transects sampled by Thompson *et al.* (2016). The purple star represents the location of the city center (Lat: 43.6561, Long: −79.3803). Inset shows the best fit regressions for the change in the frequency of HCN, *Ac*, and *Li* along an urbanization gradient, using standardized distance to the city center as a predictor. For each cline, slopes (*β*) and *P*-values for first-order (linear) terms are provided. These regression were run population-means pooled across all transects since each one showed significant clines when analyzed independently.

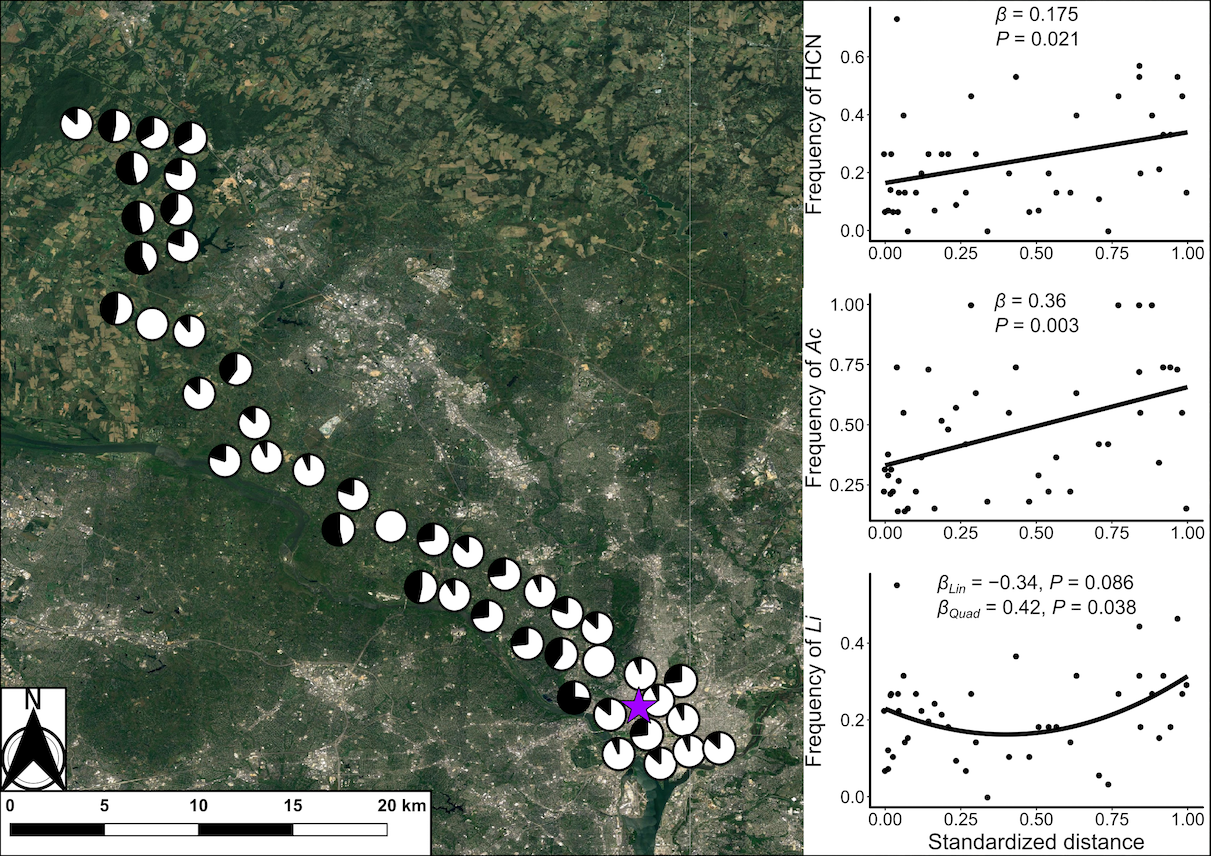

**Figure S20**: Map of the urban-rural transect for the city of Washington D. C. Populations along the transect are represented with pie charts showing the proportion of cyanogenic plants (black) in the population. Pie charts have been jittered from their actual location to improve visualization. The purple star represents the location of the city center (Lat: 38.907192, Long: −77.036873). Inset shows the best fit regressions for the change in the frequency of HCN, *Ac*, and *Li* along an urbanization gradient, using standardized distance to the city center as a predictor. For each cline, slopes (*β*) and *P*-values for first-order (linear) and second-order (quadratic, where applicable) terms are provided.
